# Gene duplication, translocation and molecular evolution of *Dmrt1* and related sex-determining genes in anurans

**DOI:** 10.1101/2025.06.11.659201

**Authors:** Sagar S. Shinde, Paris Veltsos, Wen-Juan Ma

## Abstract

Sex determination, the developmental process that directs embryos toward male or female fates, is controlled by master sex-determining genes whose origins and evolutionary dynamics remain poorly understood outside of a few model systems. In contrast to the highly differentiated sex chromosomes of mammals, birds, and *Drosophila*, most anurans (frogs and toads) maintain homomorphic sex chromosomes that exhibit a rapid turnover, even among closely related species. To uncover the mechanisms underlying the emergence of new master sex-determining genes and sex chromosome turnover, we analysed 53 published anurans and one caecilian genome (>200 Ma divergence) and available transcriptomes. We asked how often new master sex-determining genes arise by gene duplication, whether and how often gene translocation associates with sex chromosome turnover, and if new master sex-determining genes evolve under positive selection. We find that chromosome-level synteny is remarkably conserved, with only a few fusions or fissions and no evidence for translocation of four candidate master sex-determining genes (*Dmrt1, Foxl2, Bod1l, Sox3*). Only *Dmrt1* duplicated in 3 out of 50 species (excluding tetraploid *Xenopus*), and it showed strong testis-biased expression in all 8 species with available gonadal expression data. While *Dmrt1* has evolved under purifying selection, *Dmrt1* duplicates exhibit elevated nonsynonymous substitution rates and a tendency towards positive selection. Lineage-specific amino acid changes were observed in the conserved DM domain of *Dmrt1*. These results demonstrate that, in anurans, master sex-determining genes arise rarely via gene duplication, and more likely evolve via allelic diversification. Sex chromosome turnover is not associated with gene translocation, and is more likely driven by mutations on genes involved in sexual developmental pathway. All candidate sex-determining genes were under strong purifying selection, with the exception of duplications which are linked to positive selection. Our results suggest future research on anuran sex determination and sex chromosome evolution should focus on identifying allelic diversification and novel mutations on genes involved in sexual developmental pathway.

## 1. Introduction

Sex determination is a fundamental developmental process in which an embryo commits to either the female or male developmental pathway upon receiving certain cues during early embryogenesis. Despite its ubiquitous distribution across the Tree of Life, a remarkable diversity in sex-determining mechanisms have evolved^1,2^. Sex can be determined via genetic, environmental factors, or an interaction between the two^2^. For genetic sex determination, the master sex-determining gene (acting at the top of the sex-determining cascade) is located either on the Y chromosome for the male heterogametic sex chromosome system (XX females and XY males), or on the W chromosome for the female heterogametic system (ZW females and ZZ males). The XY system is characteristic of mammals, the ZW system is found in all birds, and both systems are found in insects, fishes, amphibians, reptiles and plants ^3–8^. Less well known systems are the UV sex-determining system of most algae, where sex is expressed in the haploid phase (unlike animals and plants where it is expressed in diploid cells) ^9–13^. For environmental sex determination, environmental factors or conditions influence early embryogenesis towards the female or male pathway. Environmental factors can be external, like temperature, pH, food resources, photoperiod, or biological, like social environment, or an interaction between the two, for example food allocation can be influenced by competition ^14–17^. Some organisms change sex sequentially over their lifetime, so that an adult of one sex can switch to the opposite sex depending on social conditions or age ^18^, but these will not be further discussed here. The most studied type of environmental sex determination is temperature-dependent sex determination in various turtle lineages, where the temperature at a crucial thermally sensitive embryonic stage determines sex ^19,20^. The genetic mechanisms of sex determination remain largely unknown across the Tree of Life ^1,21–23^.

For genetic sex determination, sex chromosomes have repeatedly and independently evolved numerous times across the Tree of Life, representing astonishing convergent evolution of large genomic regions^5,8,24,25^. In sharp contrast to the highly degenerated (where many functional genes are lost) sex chromosomes in mammals, most birds and *Drosophia*, sex chromosomes are largely homomorphic (appear identical with microscopy) and have minimal degeneration in most amphibians, fishes, many reptiles and flowering plants^8^. Two non-exclusive mechanisms have been proposed to explain homomorphic sex chromosome evolution, inspired by amphibian biology: i) the fountain of youth theory, which applies when males do not recombine over most of their chromosome length (including the sex chromosome), but occasional sex chromosome recombination occurs in sex-reversed XY females^26^, and ii) rapid sex chromosome turnover, where different autosomes in different closely related species quickly evolve a sex-determining role, so that there is not sufficient time for sex chromosome degeneration^27–29^. Rapid sex chromosome turnover has been identified in amphibians, fishes, reptiles and flowering plants and more are expected to be revealed with the advancing and cost effective genomic sequencing^27,29–37^. The underlying genetic mechanisms remain largely unknown, with the exception of gene translocation of master sex-determining genes in Takifugu pufferfish, salmonid fishes, and strawberries^35,37,38^.

Anurans (frogs and toads) are excellent systems to address pressing questions on the genetic mechanism of sex determination, as well as rapid sex chromosome turnover. The majority (>75%) of all studied anurans possess homomorphic sex chromosomes with little degeneration, yet extensive sex chromosome degeneration has also evolved in certain anuran lineages^39^. In addition, there is frequent and rapid sex chromosome turnover in anurans^30,31,40^, and there are documented cases both of genetic and non-genetic sex determination. Moreover, experimental manipulation during early development is easy in frogs due to their external fertilization^41,42^.

*Dmrt1, doublesex* and *mab3-related transcription factor* 1, is characterized by the presence of the DM domain, a DNA-binding motif, with an unusual cysteine-rich zinc DNA binding motif ^43^. *Dmrt1* is both indispensable for the maintenance of testis development and function, and the suppression of female-determining pathways in males across various vertebrate groups ^43–46^. Its crucial role in sex determination has been documented across vertebrates, where *Dmrt1* (or its paralog) has been identified as the master, or candidate master, sex-determining gene, including in medaka fish (*Oryzias latipes*), tropical clawed frog (*Xenopus laevis*), chicken (*Gallus gallus*) and the red-eared sliding turtle (*Trachemys scripta*), among others (reviewed in ^23^). The highly conserved role of *Dmrt1* across vertebrates highlights its fundamental importance in the genetic regulatory networks that govern sex determination and gonadal sex differentiation.

In anurans, the genetic basis of sex determination is only known for the African clawed frog *Xenopus laevis,* where a dominant, W chromosome duplication of *Dmrt1* (*DmW*) determines ovarian development^47^. Due to ancestral allopolyploidization, *X. laevis* possesses two copies of *Dmrt1*, *Dmrt1.L* (*DMRT1α*) and *Dmrt1.S* (*DMRT1β*), on autosomes 2L and 2S, respectively ^47,48^. Partial duplication of *Dmrt1.S* (*Dmrt1β*) generated *DmW*, which maintains high similarity of the 5’-coding region (exons 1-4) but lacks the 3’-coding part which includes a transactivation domain-coding region (exons 5) ^47–52^. The *Dmrt1* duplication occurred approximately 47 Mya after allotetraploidization, and *DmW* is found in several closely related *Xenopus* species. Other than *X. laevis*, *DmW* was amplified in several females but none of the males in using targeted next-generation sequencing, suggesting it is female-specific in *X. gilli*, while it is not consistently female-specific in most other *Xenopus* species^51,52^. *DmW* is required for ovary development in *X. laevis* females, and it functions as an antagonistic competitor of *Dmrt1*^50^. Both *Dmrt1* and *DmW* bind to the same regulatory regions of target genes (yet to be discovered) that are crucial in testis development. During embryonic gonad development, RNA of *DmW* was more abundant than *Dmrt1* in the primordial gonads of ZW tadpoles. This is thought to suppress the male-determining pathway because the *DmW* protein has the same binding site as the *Dmrt1* protein, but not its enhancer activity, thereby acting as a transcriptional repressor. The resulting failure of testis formation leads to ovary formation^47,50^.

Beyond *X. laevis* in Pipidae, *Dmrt1* has been identified as the candidate master sex-determining gene in Hylidae and Ranidae. In four species of the *Hyla arborea* clade, a small sex-determining region including *Dmrt1* has been identified, with polymorphism in *Dmrt1* perfectly associating with phenotypic sex^53^. In *Rana temporaria*, a homologous region has the strongest F_ST_ difference between sexes and *Dmrt1* haplotypes perfectly correlate with male testis development in two populations^41^. Beyond *Dmrt1*, the *Bod1l* gene (biorientation of chromosomes in cell division 1 like 1) was recently identified as candidate master sex-determining gene in *Bufo varilis,* because it was the only region with homozygosity in females and heterozygosity in males^54^. Two additional ‘usual suspect’ candidate sex-determining genes are *Foxl2* (forkhead box protein L2) and *Sox3* (SRY-box transcription factor 3) in anurans. *Sox3* was involved in sex chromosome system transitions (XY and ZW) in *Glandirana rugosa* and determine sex in several medaka (Oryzias) species^34,55^. *Foxl2* codes for a transcription factor essential for ovarian development and is suggested to supress testis formation, and was suggested to determine sex in tilapia and zebrafish^56,57^.

Master sex-determining genes typically evolve in two ways: gene duplication or allelic diversification. Both sex determining genes of medaka fish (*DmY*) and the African clawed frog (*DmW*) have evolved by gene duplication from *Dmrt1* ^47,58,59^. In both cases positive selection seems to have affected the duplicated genes, and in particular various amino acid mutations enhanced the binding affinity of their DM domain to DNA^60^. Beyond the two species and across anurans, it is unclear how common the mechanism of gene duplication is for the evolution of sex-determining genes. Furthermore, population genetic theory predicts that X (or Z) chromosome could play disproportionate roles in speciation and evolutionary divergence, and predicts that X- or Z-linked divergence exceeds that on autosomes (the Faster-X effect)^61,62^. New master sex-determining genes are the first to be sex-linked and are therefore predicted to evolve faster than autosomal genes^63,64^. Positive selection on newly evolved master sex-determining genes has not been tested in lineages with rapid sex chromosome turnover, where new master sex-determining genes rapidly evolve in the new sex chromosomes.

A previous study found at least 13 sex chromosome turnovers occurring within 50 Mya divergence in the 28 Ranidea true frogs. 5 chromosomes (Chr01, Chr02, Chr03, Chr05 and Chr 08 based on the reference genome of *Xenopus tropicalis*) were recruited as sex chromosomes repeatedly and in a non-random maner^30^. *Dmrt1* on Chr01 has been identified as candidate master sex determining gene in *Hyla arborea* clades and *R. temporaria,* and Chr01 has been identified to determine sex in 7 additional true frogs^30,41,53^. *Sox3* on Chr08 has been reported as candidate master sex-determining gene in *G. rugosa*^55^. Chr05 in frogs harbours *Foxl2*, also one of the most important genes in the vertebrate sex-determination cascade^65^. It codes for a transcription factor essential for ovarian development and has also been implicated in the suppression of testis formation. *Foxl2* interacts directly with *Dmrt1*, the male-determining *Dmrt1* allele blocks the expression of *Foxl2* and in turn the development of ovaries, thus producing males ^65,66^. Both gene translocation and novel mutations occur in genes involved in sexual developmental pathway are proposed to drive sex chromosome turnover in true frogs, which remain to be tested with empirical data.

In this study, we utilize published whole genome assemblies and RNAseq datasets across anurans, to address to which extent gene duplication drives master sex-determining gene evolution, as well as to which extent gene translocations drive sex chromosome turnover in anurans. We also analyse the evolution and selection affecting the ‘usual’ suspect sex-determining genes in anurans. In particular, whether the status of being (candidate) master sex-determining genes, or located on sex chromosomes, affects the selection they experience. Finally, we also analyse the conserved DM domain of *Dmrt1* across anurans, whether mutation patterns are lineage-specific and the association with functional divergence or adaptive across different frog species, and discuss how these changes might be associated with transcriptional control of downstream genes in the sex-determining pathway.

## 2. Materials and Methods

### 2.1 Anuran genomes retrieval and quality assessment

We obtained published whole genome assemblies of 53 anurans and one outgroup two-lined caecilian *Rhinatrema bivittatum* from the National Center of Biotechnology and Information (NCBI) (Table S1). Additionally, we acquired the raw genomic reads for those available on NCBI for the purpose of verification of gene duplication, as well as all available transcriptomic data (raw reads as well as transcriptomes) (Tables S2 and S3). We assessed the quality of the genome and transcriptome assemblies using Benchmarking Universal Single-Copy Orthologs (BUSCO) scores ^67^. They were either already published (Table S4A), or calculated using the Tetrapoda ortholog library (v.odb10) with the flag -m for genome or transcriptome respectively (Table S4B).

### 2.2 Phylogeny of species and DM domain sequences across anurans

We generated a phylogeny for the 54 amphibians using the available genome-wide 307 markers from Portik *et al*. (2023) ^68^. First, the available multiple sequence alignment data (54 species; Supplementary file S1) were obtained and aligned with –msaProgram in PRANK and 1000 bootstraps implemented in Guidance2. Second, the best sequence evolution model was identified using ModelTest-NG ^69^ with multiple concatenated marker sequence alignments as input. The best model --model HKY+I+G4 was selected based on the Bayesian Information Criterion score. Third, we generated maximum likelihood genome-wide trees using raxml-ng with 1000 bootstraps, employing the GTR+I+G4 model. Finally, the bootstrap-supported best tree from the raxml-ng output was visualized in FigTree (https://github.com/rambaut/figtree). However, for the phylogenetic tree of 12 anurans with chromosome-level genome assemblies and annotation, we obtained the phylogenetic tree from timetree.org and their associated divergence time in million years.

To further verify the duplicated copy was a true duplication of *Dmrt1* and not from another *Dmrt* family member, we also conducted phylogenetic analysis for the conserved DNA-binding domain (DM) of *Dmrt1,* along with DM coding sequence of all other *Dmrt* genes (i.e. *Dmrt2, Dmrt3, Dmrt4, Dmrt5*, and *Dmrt6*) across anurans. We extracted the DM domain coding sequences from all *Dmrt* genes in 53 anurans, as well as the partially duplicated region of *Dmrt1* containing the DM domain. We conducted the phylogeny using the procedure described above.

A substitution saturation test is crucial for ensuring the validity of sequence data when conducting both selection analysis and phylogenetic interpretation. We thus further assessed the sequences for substitution saturation using the index described by Xia *et al.* ^70^, implemented in DAMBE ^71^. The Index of substitution saturation (Iss) and critical Iss (Iss.c) estimate for symmetrical and extremely asymmetrical trees were reported for each group of species in our study. In all cases, we found Iss was significantly less than Iss.c, suggesting that the sequences in our study are not saturated.

### 2.3 Chromosome-level synteny analysis across anuran genomes

Twelve chromosome-level assemblies with genome annotations were used for synteny analysis (Table S4A). In addition to available genomic resources, we also considered species to provide broadly phylogenetic representation across anuran lineages. Overall on average one species from each monophyletic group was included to ensure a comprehensive comparison of synteny, with *Xenopus tropicalis* as the reference genome. Across anuran lineages with >200 million years divergence (https://timetree.org/), we performed pair-wise genome-wide synteny analysis using MCScanX ^72^. MCScanX was designed for detecting and analysing synteny and collinearity in comparative genomics, and allows to identify homologous chromosomal regions across multiple genomes or subgenomes, aligning these regions using genes as anchors. Default parameters were used to generate pairwise alignments of collinear blocks using MCScanX ^72^. The detected homologous regions across pair-wise genome comparisons were subsequently merged into a single dataset, and the synteny was visualized with TBtools-II ^73^.

### 2.4 Analyses on gene duplications of *Dmrt1*, *Foxl2*, *Sox3* and *Bod1l*

Orthologs of *Dmrt1, Foxl2, Sox3* and *Bod1l* were identified based on coding sequence similarity and micro synteny analysis (see details below). Accurate detection of orthologs and the duplicated copies was performed with a combination of high-quality genome assemblies and further validation using raw genomic reads.

For *Dmrt1*, we could directly retrieve itself and its orthologs’ coding sequences from NCBI for several species including *Xenopus tropicalis*, *Hyla sarda, Nanorana parkeri, Rana temporaria, Pseudophryne corroboree, Bufo gargarizans, Bufo bufo, Eleutherodactylus coqui,* and *Spea bombifrons*. When direct retrieval from NCBI was not possible, we performed BLASTn against the anuran genomes using the *Dmrt1* coding sequence of closely related species ^74^. BLASTn results were manually examined to confirm that the estimated length of the open reading frame (ORF) was complete. For a few species such as *Bufo gargarizans*, *Bufo bufo*, *Eleutherodactylus coqui*, and *Hyla sarda*, instead of standard 5 exons for *Dmrt1*, *Dmrt1* with 6 exons were computationally predicted in the genome annotation files, with the additional annotated exon coding for additional 14 amino acids. No clear expression data were detected from RNAseq data for this additional exon, and conserved coding sequences in all other frog lineages did not align with it. Furthermore, we examined *Rana temporaria*, utilizing published transcriptome data from different three populations and five developmental stages (G23, G27, G31, G43 and G46) to determine the number of exons expressed ^75–77^. We consistently observed the expression of five exons, which led us to focus further analysis on the conserved five exons across all species. Taken together, the extra annotated small exon of *Dmrt1* in a few anurans is most likely a bioinformatic error: gene prediction is imperfect and detailed gene structure often requires careful additional manual curation, which is especially crucial for large frog genomes (3-9.6G) ^78–80^.

We then investigated gene duplications, by identifying both whole exon and partial exon duplications, as well as complete gene duplications. These were further evaluated by analysing available genomic raw read data. All *Dmrt1* duplications, either whole gene or partial, always involved additional chromosomes. We manually inspected open reading frames (ORF) for truncations within duplicated regions, to determine whether the duplicated sequences contained premature stop codons. For the phylogenetic analysis, *Rhinatrema bivittatum* was used as the outgroup. The length of *Dmrt1* coding sequence and total gene length are described in Table S5. To inspect gene expression of both the original and duplicated copies of *Dmrt1*, we obtained RNA-seq data from those species with *Dmrt1* duplication if available on NCBI (Table S3). Furthermore, we also obtained RNA-seq data with at least one tissue from both sexes for any anuran from NCBI, which only retrieved 14 anurans (Table S6). For some of these species, genome annotations were not available, we thus used the annotated genomes of closely related species for mapping. RNA-seq reads were mapped to the corresponding genome assemblies, or the closely related reference genome, using STAR ^81^ with default parameters and performing read quantification using HTSeq-count ^82^. Similarly, we further performed gene expression analysis on the same RNA-seq data using Cufflinks ^83^. We followed the same pipeline to detect gene duplications for *Foxl2* and *Sox3*. Previous examination of *Bod1l* has shown well preserved synteny and no evidence of gene duplications across anuran genomes ^54^ so we did not analyse it here.

### 2.5 Analyses on gene translocation of *Dmrt1*, *Foxl2*, *Sox3* and *Bod1l*

To detect possible gene translocation across various chromosomes, we investigated both micro-synteny and chromosome-level macro synteny. For the *Dmrt1* micro-synteny analysis, 25 species with chromosome-level genome assemblies were used. Species with only scaffold or contig-level assemblies were excluded, as *Dmrt1* was often located on scaffolds with adjacent genes positioned separately which led to challenging synteny inference. We focused on verifying chromosome-level and gene-level synteny for *Dmrt1* and its five flanking genes on both upstream and downstream regions. Of the 25 species considered, genome annotations were available for 12 species, which were used for the genome-wide synteny analysis. Despite the availability of annotations, some species lacked complete annotations for the syntenic genes. In such cases, we performed BLAST search to verify gene presence. If a gene was not assembled, no strong blast hit was detected, or the gene was entirely missing from the genome. Thirteen genomes lacked annotation data. To validate synteny for these chromosome-level assemblies, we performed BLAST search using annotated gene sequences from other species available on NCBI. While the exact start and end positions were not always determined, we identified the highest BLAST hit and used it to infer gene location. We used the same pipeline to conduct micro-synteny analysis for *Bod1l1*, *Sox3*, and *Foxl2.* Finally, the chromosome-level macro synteny was conducted as part of genome-wide synteny, see details in the earlier section (2.3 Chromosome-level synteny analysis across anuran genomes).

### 2.6 Molecular evolution and selection analysis for *Dmrt1*, *Foxl2* and *Sox3*

To investigate molecular evolution and selection type acting on candidate sex determining genes, we used three branch-specific models implemented in PAML using the codeml program ^84,85^. Selection inference was based on the *d_N_/d_S_* ratio (*ω*), following a maximum likelihood approach. We tested three branch-specific models M0 (null model) assuming a single *ω* for all species, branch-neutral (alternative model) assumes the foreground species evolves under neutral evolution, and branch-free (alternative model) estimates *ω* independently for the foreground and background species. The analysis included *Dmrt1* coding sequences and its paralogs from 54 species, including the duplicated *DmW* of *Dmrt1* in *X. laevis* and the duplicated *Dmrt1* copy *in Pelodytes ibericus*. The genome-wide phylogenetic tree was used for such analysis. The codeml program was executed using F1×4 and F3×4 codon frequency models to assess the signatures of relaxation, or intensification of purifying selection, or positive selection.

Additionally, to evaluate the impact of duplication of *Dmrt1* in *Pelodytes ibericus*, we compared the results with and without the duplication using the branch-test model. To further assess *ω* patterns, we calculated *ω* using the free-ratio model in codeml, which estimates *ω* independently for each branch. The *ω* values of the foreground branch from the branch-test model was compared with the results from the free-ratio *ω* analysis, which remained largely identical. Lastly, we used the PAML branch-site model to identify specific codon sites under positive selection. Finally, similar branch test analyses were performed for the other sex-determining genes *Foxl2*, *Sox3*, as well as one randomly selected transcription factor gene E2F transcription factor 1 (E2F1) for a subset of 12 species.

### 2.7 Evolution of conserved DM domain in *Dmrt1*

We aligned the amino-acid sequences from 53 anuran species using PRANK ^86^, applying default settings. To evaluate alignment reliability, we performed 1000 bootstrap replicates with the PRANK ---iterate option. All sequences were inspected to ensure they begin with the initiating methionine (M) residue. The final, consensus alignment was visualized in Jalview (alignment viewer) ^87^. We then mapped unique and clade-specific amino-acid substitutions onto the alignment and corresponding phylogenetic tree, manually verifying each site against the phylogeny to confirm its consistency and taxonomic specificity. The *Dmrt1* protein sequence similarity score was calculated and visualized for the whole *Dmrt1* gene as well as the DM domain using Jalview.

## 3. Results

### 3.1 Strong preservation of chromosome-level synteny across anuran genomes

We obtained 54 amphibian genomes from NCBI, 53 anurans and 1 two-lined caecilian (*Rhinatrema bivittatum*) which serves as an outgroup (Table S1). Using *Xenopus tropicalis* as the reference genome, we inferred chromosome-level synteny, gene translocations and duplications across the anuran genomes. All available (12) chromosome-level assemblies with genome annotations were used for synteny analysis. Assembly completeness was assessed using BUSCO with the tetrapoda_odb10 dataset (n=5310) and resulting scores were >85%, with 2 exceptions (around 75%) (Table S4). The phylogenetic tree across the 12 anurans was generated from timetree.org, with estimated divergence time in million years (Figure 1a).

**Figure 1.**
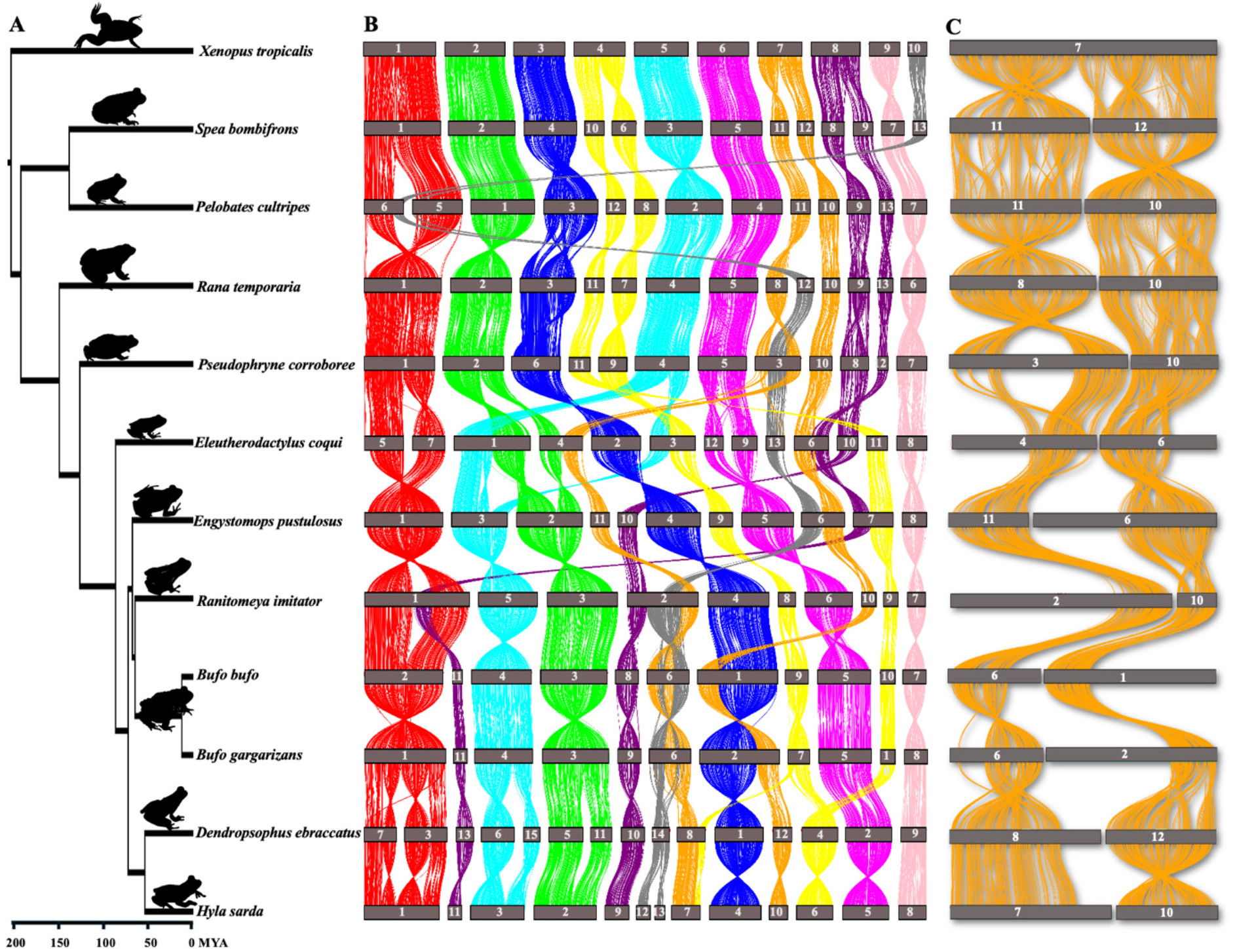
Genome wide synteny across 12 anuran genomes with chromosome-level genome assemblies and genome annotation. A). The phylogenetic tree from timetree.org. B). Chromosome level synteny. Chromosomes are numbered and ordered by descending size (chromosome 1 is the largest, chromosome 14 is the smallest). C). Example of chromosome 7 in *X. tropicalis* which is involved in sex determination in both *X. tropicalis* and *Hyla sarda* ^88,89^, shows it has undergone fissions and fusions in various anuran lineages.

Genome-wide pairwise synteny was assessed based on sequence alignments and collinear blocks were generated using MCScanX and visualized in TBtools-II. The order of each pairwise comparison followed the smallest phylogenetic distance, starting with the reference genome *X. tropicalis* (Figure 1a). The chromosome-level synteny was well conserved across all anuran lineages, spanning >200 Mya divergence (Figure 1a). While there are several chromosome fissions and fusions in various frog lineages compared to the karyotype n=10 of *X. tropicalis,* synteny blocks between these fission/fusion chromosomes were conserved (Figure 1b). For instance, *X. tropicalis* chromosome 7 is split into chromosomes 11 and 12 in *Spea bombifrons*, chromosomes 10 and 11 in *Pelobates cultripes*, chromosomes 8 and 10 in *Rana temporaria*, chromosomes 3 and 10 in *Pseudophrynes coorroborea*, partial chromosome 4 and 6 in *Eleutherodactylus coqui*, partial chromosome 6 and chromosome 11 in *Engystomops pustulosus*, partial chromosome 2 and chromosome 10 in *Ranitomeya imitator*, partial chromosome 1 and 6 in *Bufo bufo*, partial chromosome 2 and 6 in *B. gargarizans*, chromosome 8 and 12 in *Dendeopsophus ebraccatus*, as well as chromosome 7 and 10 in *Hyla sarda* (Figure 1c). Similarly, the synteny of *X. tropicalis* chromosome 1 corresponds to chromosome 5 and 6 in *Pelobates cultripes,* or chromosome 5 and 7 in *Eleutherodactylus coqui*, or chromosome 3 and 7 in *Dendropsophus ebraccatus* (Figure 1b), suggesting independent chromosome fissions took place in various frog lineages. Large inversions were rare across highly diverged anuran genomes. The noticeable ones are 2 big inversions on chromosome 1 of *Hyla sarda*, corresponding to chromosome 3 and 7 of *Dendropsophus ebraccatus* (Figure 1b), suggesting the inversions’ involvement in the chromosomal fission in this species. Similarly, we identified local inversions across multiple chromosomes, such as chromosome 4 and 5 between *B. bufo* and *B. gargarizans*, and chromosome 11 between *Spea bombifrons* and *Pelobates cultripes* (Figure 1b, 1c).

### 3.2 *Dmrt1* rarely duplicated in various anuran lineages with fully sequenced genomes

Master sex-determining genes can evolve via gene duplication or allelic diversification. We first investigated whether *Dmrt1* was recruited as master sex-determining gene via gene duplication in the anuran lineages with available whole genome sequences (Table S1). In addition to BLAST (based on coding sequencing of *Dmrt1*) search, we also performed phylogenetic analyses of the DM domain across all *Dmrt* genes and their detected duplicated copies, to provide a robust and thorough verification of all *Dmrt1* duplications across various anuran genomes. For the phylogeny of the DM domain, all duplicated copies largely clustered with *Dmrt1*, supporting their origin from *Dmrt1* duplication (Figure S1).

Across the 53 anurans, 3 are tetraploid (*X. laevis, X. petersii,* and *X. borealis*) which had 2 *Dmrt1* copies that arose via whole genome duplication. Additionally, *DmW* arose from duplication of *Dmrt1.S* (another copy is *Dmrt1.L*) and was recruited as master sex-determining gene in *X. laevis* but was not consistently amplified in females in other two species^51,52^, which questioned *DmW*’s role as sex-determining gene in two other *Xenopus* species. For the remaining 50 species, there are only 3 cases of near-complete gene duplication involving 3 (out of 5) or more exons of *Dmrt1*, and 5 cases of duplication of only 1 or 2 *Dmrt1* exons, scattered across the phylogeny (Figure 2). All duplications were verified using raw genomic sequencing reads (Table S2). Since the outgroup and ancestral branch of anuran lineages all have a single copy of *Dmrt1*, *Dmrt1* has probably undergone approximately 8 independent duplication events (Table S7).

**Figure 2.**
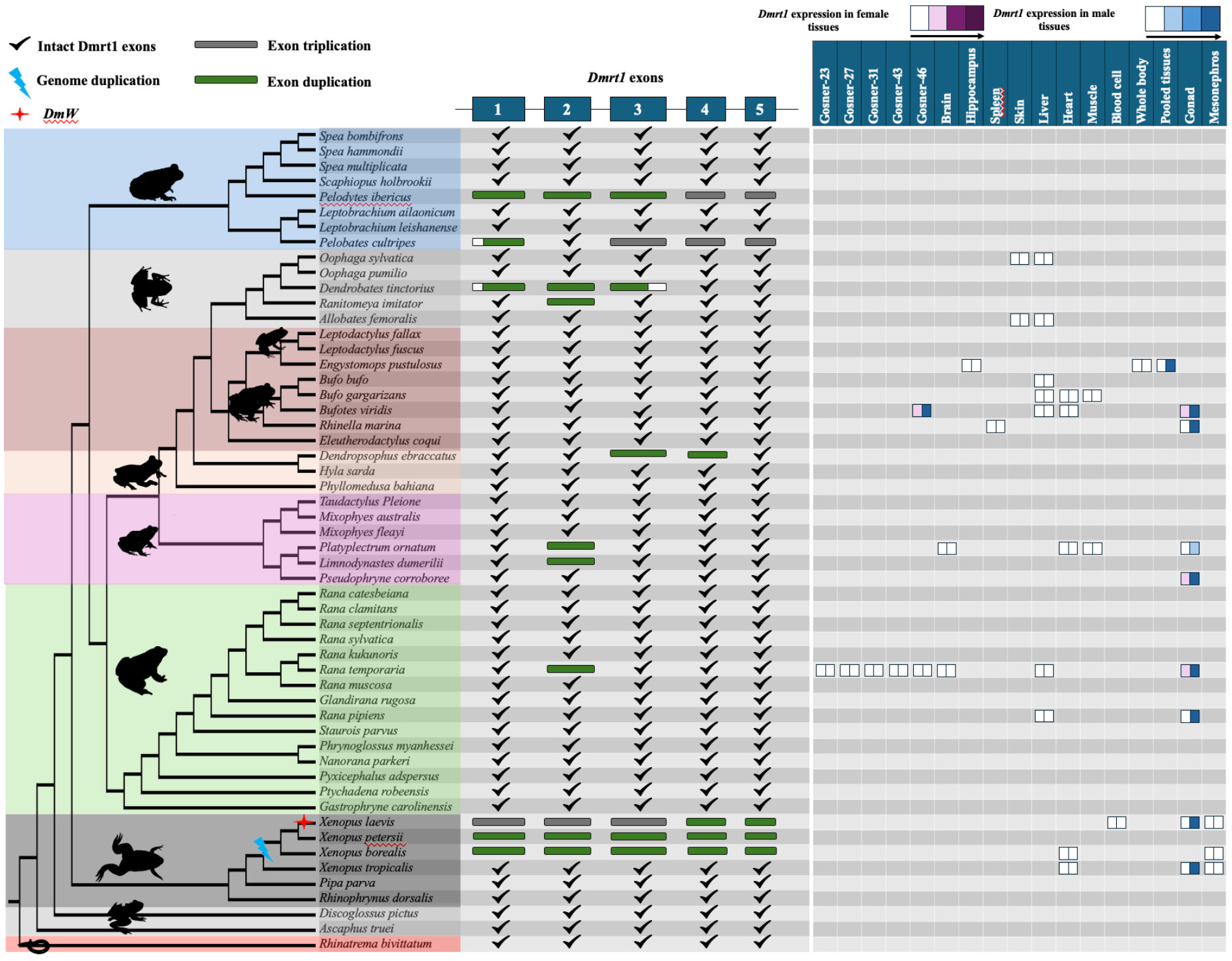
*Dmrt1* coding sequence structure and detected duplicated copies across 54 amphibia. The phylogenetic tree was conducted with multiple-loci alignment from Portik *et al.* (2023) ^68^. *Dmrt1* exon duplications are colour coded (2 copies in green, 3 copies in grey). *Dmrt1* expression is plotted when data from both sexes from at least one tissue are available. The color gradient indicates expression in female (red) and male (blue) tissues. Colour intensity indicates the scale of gene expression, the four expression bins are < 100 reads,100 – 500 reads, 500-2000 reads, >2000 reads.

Near-complete duplication of *Dmrt1* was detected in 3 species (Figure 2). First, in *Pelodytes ibericus*, *Dmrt1* localised in the syntenic region on homologous chromosome 1 (of *X. tropicalis* reference genome), a complete duplicated copy resided on chromosome 10 and a third copy composed of only exons 4 and 5 was found on chromosome 6. Haplotype validation of the duplicated sequences was not possible due to lack of raw genomic reads. For the complete duplicated copy, the intact open reading frame was 305 amino acids, however, no transcriptomic data was available to detect gene expression level. Second, one near-complete duplication of *Dmrt1* was identified in *Pelobates cultripes* (missing exon 2), and an additional third copy composed of only exons 3, 4 and 5. All three copies, each with a distinct haplotype, were further supported by raw genomic reads (Table S2). The intact *Dmrt1* gene was located on chromosome 1, with the near-complete duplicated copy on chromosome 11 and the third copy on chromosome 14. The function of the truncated duplicated copies is unclear. While no premature stop codons were detected in both duplicated copies, there was no expression of either the near-intact or duplicated *Dmrt1* copies in tadpoles and adult tissues (Figure S2-S4). The conserved DM domain is primarily located on exon 1 and beginning of exon 2, which are missing from the truncated duplicated copies. Third, in *Dendrobates tinctorius*, the duplicated copy composed of a complete exon 2 and parts of exons 1 and 3. The two *Dmrt1* copies located on two different scaffolds, and could be distinguished from haplotype analysis using raw read data. Similarly, while there were no premature stop codons in the duplicated copy, transcriptomic data of skin, liver, gut, and brain tissues did not show it (or *Dmrt1*) *to* be expressed (Figure S5, S6).

Finally, approximately 4 independent duplications of one or two exons of *Dmrt1* have been detected in 5 frog species (Figure 2). Among them, is the European common frog *Rana temporaria*, one of the most studied anurans where the *Dmrt1* gene is located within a syntenic region on chromosome 1 and has been identified as a candidate master sex-determining gene ^41,42,90^. A complete copy of exon 2 was duplicated on chromosome 10, which was further supported by identification of two haplotypes using raw sequencing reads. Transcriptomic analysis showed a strong testis-specific expression, no expression in brain and liver and very low expression in different developmental stages (Gosner stage 23, G27, G31, G43, and G46) (Figure 2) (Figure S7, Table S6). However, the duplicated exon 2 is not expressed in any tissue (Figure S8). In other two species (*Platyplectrum ornatum, Limnodynastes dumerilii*) with duplicated copy of only exon 2, both copies were located on two different scaffolds. Due to low genome assembly quality, it remains unclear whether two copies are located on the same chromosome. We detected two different haplotypes for exon 2 in both species using the raw sequencing reads. In *Platyplectrum ornatum, Dmrt1* was extremely biased in male testis with very low expression in ovary, and was not expressed in somatic tissues (Figure 2). It is possible that the duplication of exon 2 may have occurred in a common ancestor of the two species.

In *Dendropsophus ebraccatus*, a duplication of exons 3 and 4 was identified. The intact exons are located on chromosome 3 within a syntenic region, while the duplicated exons are on chromosome 1, positioning close to each other. The assembled and annotated genome reveals that the duplicated copy of *Dmrt1*is located on a different chromosome compared to the syntenic copy of the gene. Both copies of *Dmrt1* gene are identical, with no haplotype differences observed in the supporting genomic raw reads as well as in the assembled genome, suggesting a recent duplication event with minimal sequence variation. Transcriptome data from whole-body tissue confirmed high expression of *Dmrt1* (Figure S9). In *Ranitomeya imitator*, a duplication of exon 2 in *Dmrt1* was identified on chromosome 3, with the intact gene located on chromosome 1 within a conserved syntenic region. Two haplotypes were observed in the genome, but genomic raw data is not currently available for further verification. Transcriptome data from skin and brain tissues showed no expression of *Dmrt1* and duplicated region in this species (Figure S10, S11)

Based on differential expression data from 14 species, *Dmrt1* always shows strong expression in testes (Figure 2), whereas its expression in ovaries is minimal or absent. There is no detectable expression of *Dmrt1* in various somatic tissues of either sex, nor during the developmental stages until G46 (froglet), underscoring its specialised role in the reproductive system (Figure 2, Figure S12, S13, Table S6). The near testis-specific expression of *Dmrt1*, aligns with the well-established role of *Dmrt1* as a critical regulator of testis development and male sex determination. The conserved nature of this expression profile across phylogenetically distant species highlights the evolutionary importance of *Dmrt1* in male reproductive differentiation.

We also investigated gene duplication for two other strong candidate sex-determining genes in frogs, *Sox3* and *Foxl2* and found no evidence for it. One possible duplication of *Foxl2* in *Rana muscosa* is likely an assembly error as there is no sequence divergence between the two copies, and no excessive genomic reads associated with the gene. Furthermore, *Bod1l* also has no duplication across diverged anuran genomes^54^. Taken together, the current evidence suggests that *X. laevis* is a rare exception to the rule and master sex-determining genes do not commonly evolve in anurans via gene duplication.

### 3.3 No evidence for gene translocation of key frog sex-determining genes driving sex chromosome turnover in frogs

We assessed whether gene translocation of the ‘usual suspect’ master sex-determining genes, *Dmrt1, Sox3, Foxl2 and Bol1l* drives sex chromosome turnover in frogs. We investigated both micro-synteny (consisting of 5 upstream and 5 downstream genes) and macro-synteny at the chromosome level across anuran genomes. For *Dmrt1*, the genomic regions flanking *Dmrt1* exhibit highly conserved synteny (Figure 3 & 4, Table S8) within 12 anurans. Using the gene order from the reference genome of *X. tropicalis*, *Dmrt3*, *Dmrt2*, *Smarca2*, *Vldlr*, and *Kcnv2* were identified upstream, while *Kank1*, *Dock8*, *Cbwd3*, *Cytbp4502j6-like*, and *FoxD5a* were downstream (Figure 3A, Table S8). Various *Cytb* genes were annotated surrounding *Dmrt1* across anuran genomes (Figure 3A). In certain anurans, species-specific uncharacterized *Loc* genes have been assembled in the syntenic (Figure 3A, Table S8). In the outgroup *Rhinatrema bivittatum*, the upstream region contains five conserved genes. On the downstream region, the first three genes adjacent to *Dmrt1* (*Kank1*, *Dock8*, *Cbwd3*) are conserved, but the other two (*Cytbp4502j6-like* and D4 dopamine receptor-like genes) localise on different chromosomes. Instead, *Pgm5* and *Tmem252* are present. Overall, the *Dmrt1* micro-synteny inferred via a BLAST approach was largely conserved in 25 species with chromosome-level assemblies, and no local inversions were detected (Figure 3A & 4). For other anuran genomes (28) with fragmented assemblies, while these *Dmrt1* flanking genes are generally located near each other, the fragmented assemblies do not allow to detect translocations within the synteny block.

**Figure 3.**
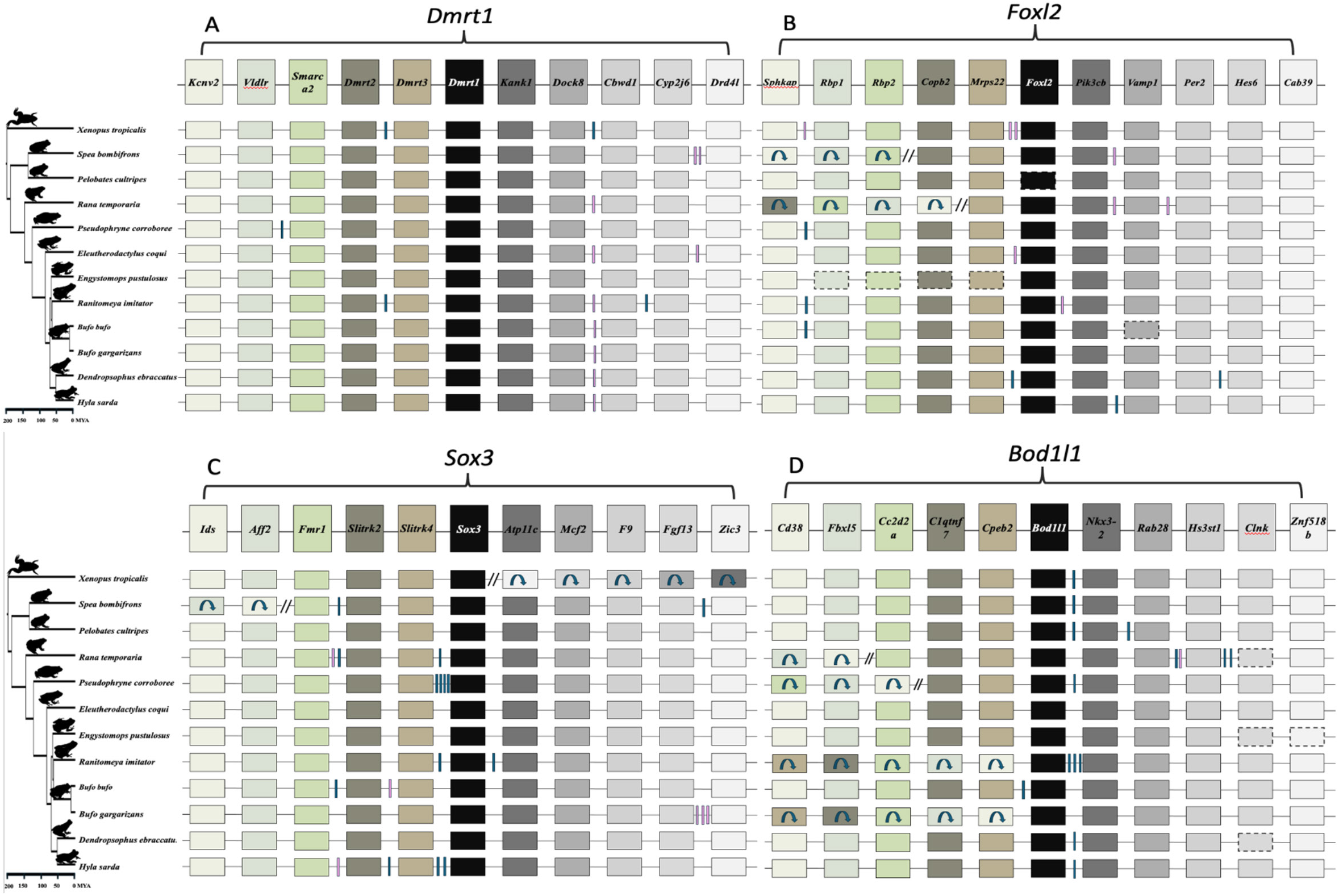
Gene order of the four candidate master sex-determining genes, (A) *Dmrt1*, (B) *Foxl2*, (C) *Sox3*, and (D) *Bod1l* across 12 anurans with chromosome genome assemblies. The top row shows the focal gene, its immediate upstream and downstream 5 genes based on the reference genome of *X. tropicalis*. Boxes with arrows indicate gene inversions, while paired tilted lines represent multiple genes with inverted orientations. Vertical blue boxes mark unique, uncharacterised LOC genes that are species-specific, and pink lines denote other species-specific genes respectively. Genes missing from the genome assembly are enclosed in dotted boxes. Detailed list of inversions and species-specific gene list can be found in Table S8-S11.

Similar analysis of micro-synteny was conducted for *Foxl2, Sox3* and *Bodl1l*, and it was also well conserved (Figure 3B, 3C, 3D, Table S9-S11). In *Foxl2*, the overall gene order remained intact on both sides, except in *Rana temporaria*, where a local inversion on the downstream region, following the *Mrps22* gene, caused the last four genes to be rearranged in the syntenic region (Figure 3B & 4, Table S9). For *Sox3*, in *X. tropicalis*, a local inversion on the upstream region caused the rearrangement of all 5 genes after *Sox3*, while the downstream region remained intact (Figure 3C). Similarly, in *Spea bombifrons*, the last two syntenic genes on the downstream region were subject to local inversion, leading to a change in gene order. No local inversions were observed in other species for *Sox3* (Figure 3C & 4, Table S10). In addition, for *Bod1l1* gene synteny, local inversions resulting in changes in gene order were detected on the downstream region but the upstream region remained intact (Figure 3D, Table S11). The chromosome level macro-synteny of all ‘usual suspect’ candidate sex-determining genes was largely preserved across the various anuran genomes (Figure 1B & 4).

A previous study found at least 13 sex chromosome turnovers in 28 Ranidea true frogs^30^, and suggested that it could be driven by gene translocation, and/or novel sex-determining genes in the sexual development pathway. *Dmrt1* and its paralogs have been suggested as candidate sex-determining genes in 3 frog species including *X. laevis*, *R. temporaria*, *H. arborea*, and chromosome 1 on which *Dmrt1* was identified as the sex chromosome in 8 true frog species ^30^. Both *Dmrt1* micro-synteny and chromosome 1 macro synteny were strongly preserved in all 25 anurans with chromosome-level assemblies spanning >200 Mya divergence (Figure 4, Table S8), suggesting that *Dmrt1* translocation is unlikely to have driven sex chromosome turnover in Ranidae, as well as across various anuran lineages. A similar result for *Foxl2*, *Sox3* and *Bol1l*, suggests that repeated gene translocation is not the general genetic mechanism driving sex chromosome turnover across anurans.

**Figure 4.**
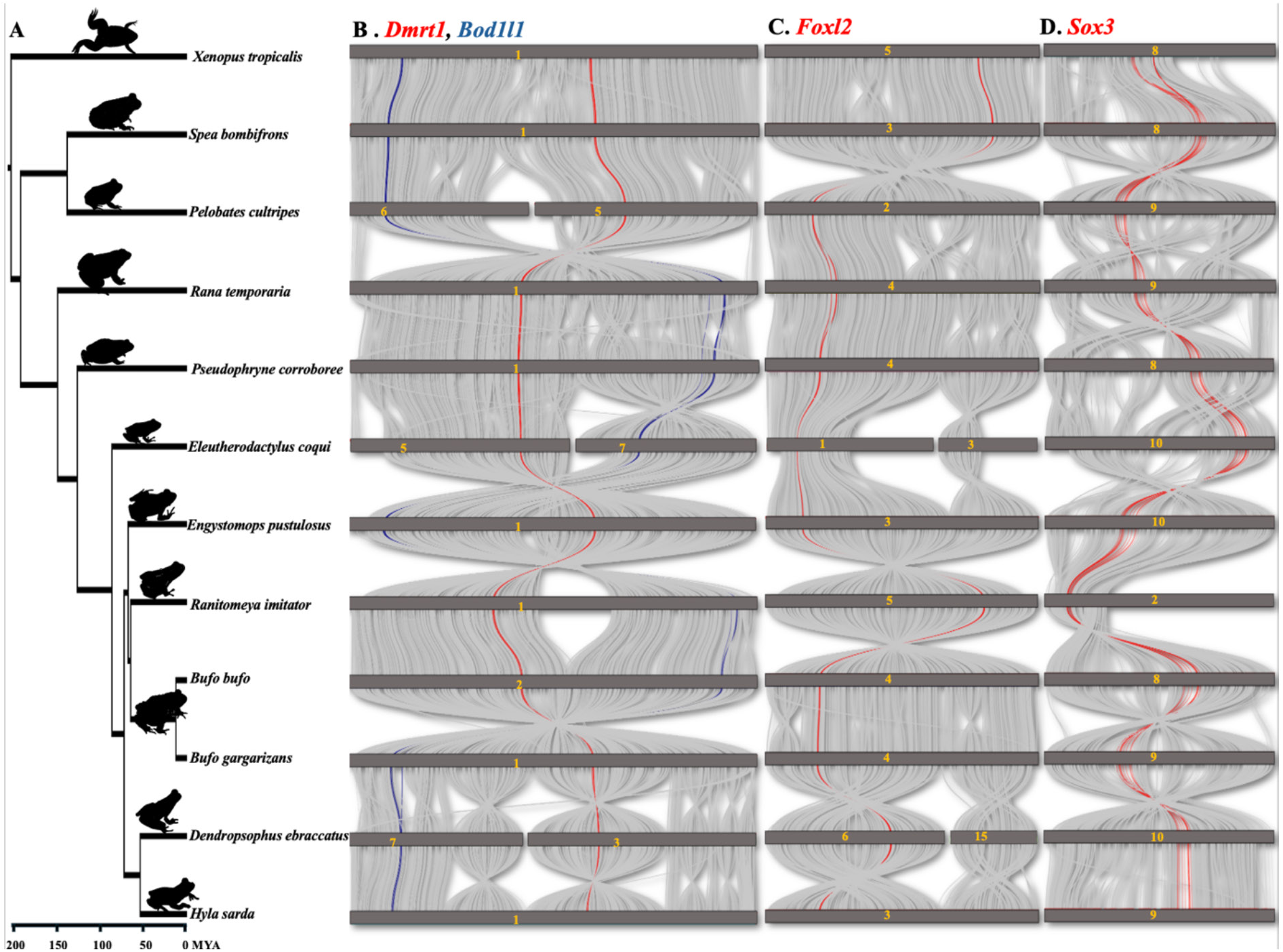
A) The phylogenetic tree of 12 studied anurans alongside graphs of chromosome level synteny for the micro-synteny region of B) *Dmrt1* and *Bod1l1,* C) *Foxl2* and D) *Sox3*.

### 3.4 Strong purifying selection acting on *Dmrt1* and other sex-determining genes

We investigated the molecular evolution of candidate master sex-determining genes *Dmrt1*, *Foxl2*, *Sox3*, *Bod1l* and their paralogs. The free ratio models, implemented in PAML, were used to infer omega (*ω,* d_N_/d_S_) which is the ratio of non-synonymous (*d_N_*) to synonymous (*d_S_*) substitutions across 53 anurans (Figure 5, Figure S14). If *ω* < 1 and smaller than background species, it suggests purifying selection. If *ω* = 1 the sequences evolve neutrally, while if *ω* > 1 the sequences may have evolved under positive selection. We first tested the global patterns of d_N_/d_S_ along all branches of the phylogenetic tree (Figure S14). We identified two species, *Oophaga_pumilio* and *Rana septentrionalis*, where *d_N_* exceeded *d_S_*, resulting in *ω* > 1. However, this is unlikely the result of positive selection because there were very small d_N_ (0.003556 and 0.007135) and even smaller d_S_ values (0.000004 and 0.000007) (Table S12A-B). For the vast majority of anurans, *d_N_* was 0-0.05 and *d_S_* was 0.05-0.4, indicating purifying selection (Figure 5, Table S12A-B). In the case of the master sex-determining gene *DmW*, *ω* is 0.47, suggesting reduced purifying selection and tendency towards positive selection, while the autosomal (original) copy *Dmrt1* with a *ω* of 0.05 was under strong purifying selection.

**Figure 5.**
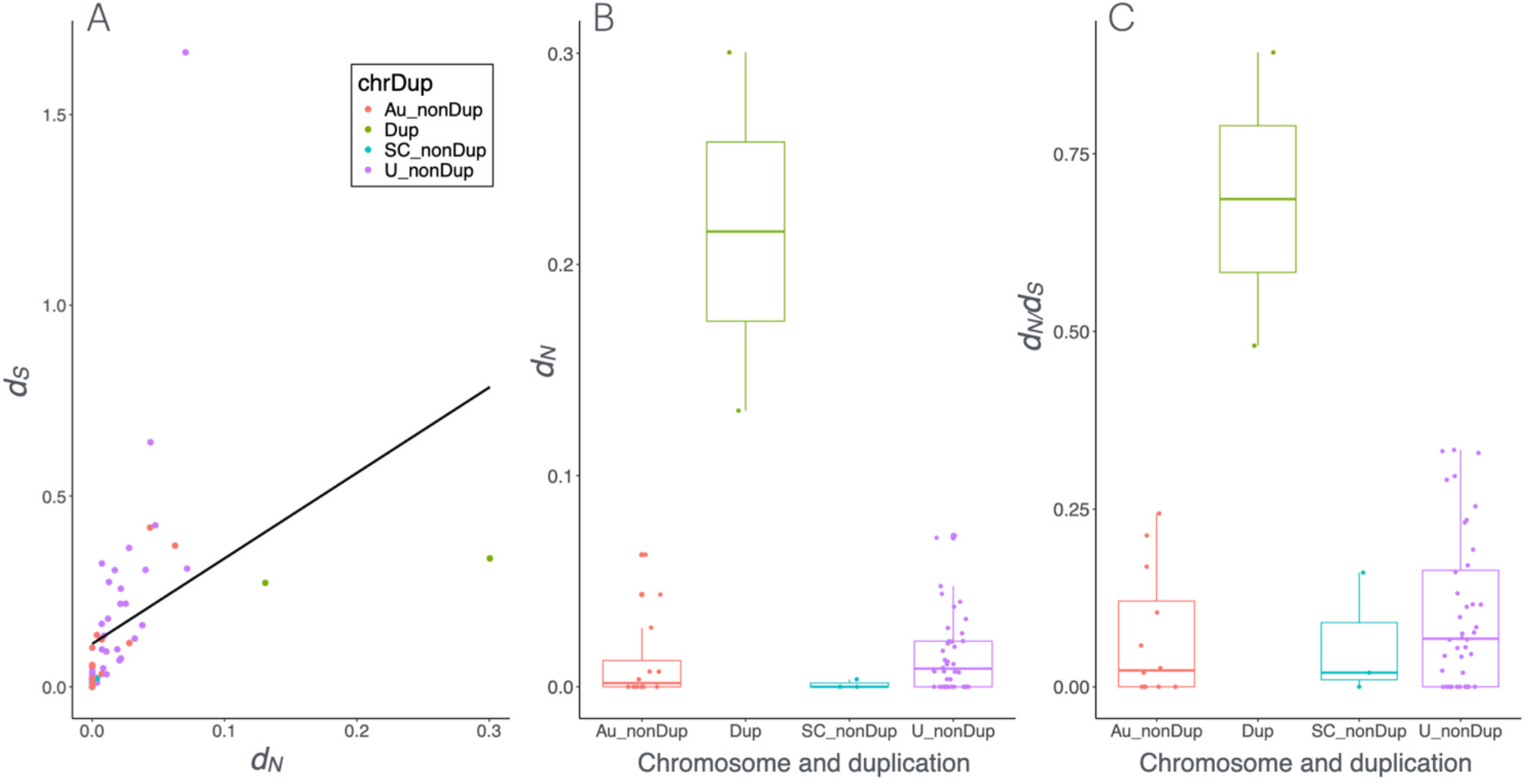
Plot of *d_N_* and *d_S_* values of *Dmrt1* and its duplicated copies for 53 anurans, including the status of sex chromosomes reported in the literature among 53 anurans. A) Spearman’s correlation between *d_N_* and *d_S_* values. B) Boxplot of *d_N_* values C) boxplot of *d_S_* values across duplication and non-duplication status, for the latter also divided in autosomal, sex chromosome and unknown. chrDup: chromosome and duplication status, Au: autosome, SC: sex chromosome, U: unknown, nonDUP: non-duplicated copy, Dup: duplication.

We then performed a branch model test on all 53 anurans, including *DmW* and both duplicated copies of *Dmrt1* from three *Xenopus* species (*Dmrt1L* and *Dmrt1S*). By comparing *ω* of the test species to that of the 52 background species, we detected varying selection pressures on *Dmrt1* or its paralogs. Among the test species, the ω of 17 species was greater than the background species and less than 1, suggesting reduced purifying selection. Statistically significant results were found in two species, the duplicated copy of *Pelodytes ibericus* and *DmW* in *X. laevis* (Figure 5A, 5B, Table S12C). In contrast, ω of *Rana septentrionalis* and *Oophaga pumilio* was greater than the background species and greater than 1, implying positive selection. However, this was not statistically significant, and likely the result of very low *d_N_* and even lower *d_S_* values for these species (Table S12C). For the remaining 37 species, ω was lower in the test than the background species and smaller than 1, indicating purifying selection (Figure 5A). Only *Discoglossus pictus, X. tropicalis* and the outgroup *Rhinatrema bivittatum* showed statistically significant intensification of purifying selection (Table S12C). ω of the duplicated copy of *Pelodytes ibericus* is 0.92, appears to be under strong tendency towards positive selection.

We further used the branch-site model to identify which codon sites were under positive selection and assessed the distribution of sites in different selection schemes. Overall, more than 39 species showed that approximately more than 70% of sites were under purifying selection, while the remaining sites were under neutral selection. Only 14 species showed evidence of a few sites with positive selection with or without significance (Table S12D). Interestingly, *DmW* in *X. laevis* and the duplicated copy of *P. ibericus* showed multiple sites under positive section (Table S12D).

To understand the relationship between selection type and function of *Dmrt1*’s role as master sex determination or sex linkage, we applied generalized linear models on all *d_N_, d_S_* values in the branch model. Overall, the vast majority of species showed very low values of *d_N_* and slightly higher values of *d_S_*, the values were significantly positively correlated (Figure 5A, Spearman’s correlation, *ρ* = 0.74, *P* < 0.001), suggesting strong purifying selection. Furthermore, sex chromosome status does not affect the selection but gene duplication significantly affects ω (Glm: ω ∼ chromosome + duplication, family = gamma, for sex chromosome: *P* = 0.27; for duplication, *P* = 0.002, Figure 5A). In particular, *d_N_*, *d_S_* and ω values were significantly higher for *Dmrt1* duplicated copies (*DmW* in *X. laevis* and the duplicated copy in *P. ibericus*, Figure 5B & 5C, Figure S15). For instance, in *Rana temporaria, Bufotes viridis*, *Rana clamitans*, *R. catesbeiana, and R. kukunoris,* when *Dmrt1* is located on sex chromosomes, the gene was under strong purifying selection but not significantly different than those on autosomes. Taken together, *Dmrt1* is under strong selective constraint overall. This could be due to its crucial role in testis formation, in meiosis ^91^, involvement of ovary function^92,93^, or a combination of these ^23^.

Similarly, for the two other candidate sex-determining genes, *Foxl2* and *Sox3*, *d_N_/d_S_* analyses across anurans revealed that both *d_N_* and *d_S_* were small and *d_S_* were consistently higher than *d_N_* (Table S13A, S14A). In terms of selection patterns, out of 53 analysed anurans, more than 32 species exhibited strong purifying selection on both *Foxl2* and *Sox3* (Table S13B & S14B). In the remaining species, a reduction in the strength of purifying selection was observed. While the variation in *ω* between background and foreground branches was generally minor, some species showed substantial differences, indicating a relaxation of purifying selection. Given *Foxl2* and *Sox3* have not been clearly identified as candidate sex-determining genes across anurans, so glm analysis on sex chromosome status was not conducted. Taken together, these results suggest that major candidate sex-determining genes are predominantly evolving under purifying selection across anurans, with occasional relaxation events in certain lineages.

### 3.5 Conservation and lineage-specific mutations on the DM domain

The conserved DM domain of *Dmrt1* consists of 54 amino acids, spanning positions 29 to 82 of *Dmrt1* protein on exon 1 and 2 (Figure 6). The DM domain is one of the most conserved motifs, reflected by the very high protein sequence similarity (0-1 with 1 is 100% similarity), with on average 0.96 score comparing to whole *Dmrt1* average score of 0.83 (Figure 6, Figure S16, S17). Despite the conserved DNA-binding function, we observed lineage-specific amino acid substitutions that suggest potential functional divergence or adaptation. First, within the DM domain, we identified Ranidae lineage-specific amino acid substitutions combination uniquely in 14 species of this lineage. These specific amino acid substitutions combination include: an Leucine (L) to Methionine (M) change at position 30, an Alanine (A) to Serine (S) change at position 34, a Proline (P) to Leucine (L) at position 44, and a D (aspartic acid) to E (Glutamic Acid) change at position 56 in nearly all 14 species (Figure 6, yellow highlight). Among these amino acid sites, codon site selection analysis suggests mutations were not significantly under positive selection (Table S12D). Beyond the Ranidae group, four additional species *Oophaga pumilio, Oophaga sylvatica, Dendropsophus ebraccatus,* and *Pipa parva* also exhibit the D to E substitution at position 56, suggesting a convergent evolution at this position. This convergence may reflect selective pressure on this residue. Possibly these specific mutations modify the DNA-binding affinity or protein-protein interactions of the *Dmrt1* protein. The recurrence of these mutations specifically within Ranidae suggests possible lineage-specific functional tuning of *Dmrt1*, what the exact function of these substitutions combination, why this occurs in Ranidae are interesting and require further investigation.

**Figure 6.**
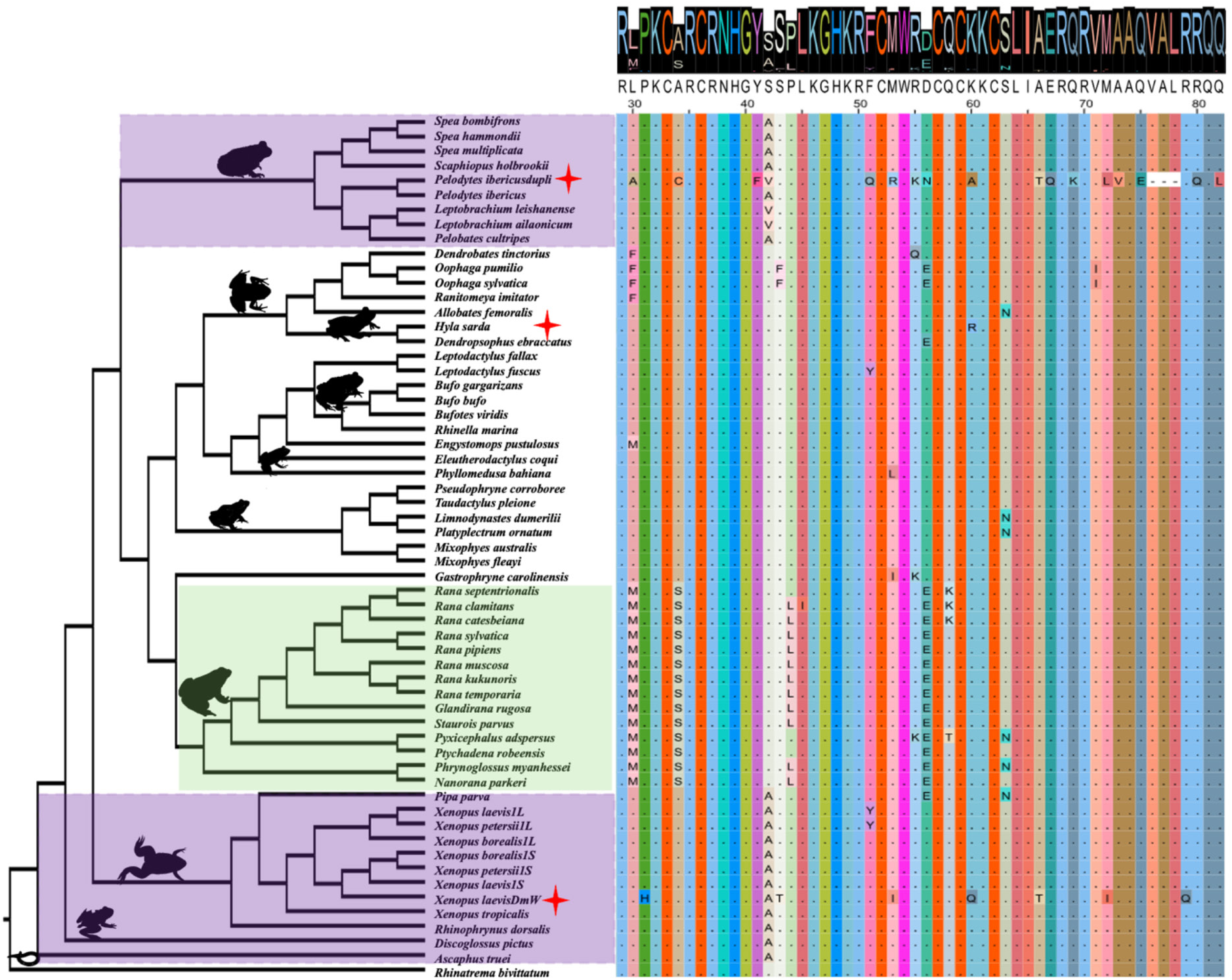
The protein sequence alignment and protein similarity analysis for the DM domain of *Dmrt1* across 53 anurans. The colours correspond to amino acid identity, and the amino acid letter size correspond to the sequence similarity across anurans. The amino acid sequencing similarity was scored with an average of 0.96 (out of 1 as 100% similarity without mutation), indicating high sequence similarity. Yellow shading highlights the Ranidae lineage with lineage-specific amino acid mutation combination, and purple shading possibly indicates retaining of ancestral amino acid in the basal lineages at position 42. Red star indicates certain amino acid mutation sites are under positive selection in three species (position 64 in *H. sarda*, and various sites in two *Dmrt1* duplicated copies, *DmW* of *X. laevis* and *Pelodytes ibericus*).

Another interesting finding is the at position 42 (purple highlight), 17 species at ancestral branches harbour an A (Alanine) amino acid, two species have a Valine (V), and the remaining species have a change from Alanine (A) to Serine (S). The parsimonious scenarios are: i) one mutation from S to A took place once in the ancestral branches, and 3 species had another round of 2 independent mutation events from S to V. This requires 2 mutation changes; ii) in the derived lineages one mutation from A to S took place, and 3 species had another found of 2 independent mutation events from A to V. Both scenarios require similar mutation changes, and it is unclear which event is most likely. Nevertheless, selection analysis on codon site showed no significant for positive selection at this position in all anurans. The repeated occurrence of the change suggests that certain sites within the DM domain may be subject to change, which could reflect on the capability of DNA binding affinity and affect various pathways it interacts with.

Finally, in *H. sarda* (red star), we identified a site under positive selection in the DM domain at position 60, with an amino acid change from Lysine (K) to arginine (R). Multiple sites (positions 43, 53, and 72) of the *Dmrt1* duplicated copy *DmW* in *X. laevis,* and the *Dmrt1* duplicated copy of *Pelodytes ibericus*, are under positive selection, although these are not statistically significant (Table S12D).

## 4. Discussion

Comparative genomic analysis revealed a highly conserved chromosome-level synteny across anuran genomes spanning >200 million years divergence, with certain chromosomal fusions and fissions yet also preserving good collinearity. We further detected rare gene duplications of *Dmrt1* (3 out 50, excluding tetraploid *Xenopus* frogs), and no duplication was detected for *Foxl2*, *Bod1l* and *Sox3*. *Dmrt1* showed a near testis-specific expression across all 8 anurans with gonadal expression data. No gene translocation events were detected across anuran genomes for all 4 sex-determining genes. Selection analyses showed 4 genes were under purifying selection. For *Dmrt1*, the status of master sex-determining gene or sex linkage, did not affect the selection scheme, but gene duplications significantly contributed to positive selection. Finally, at the conserved DM domain region of *Dmrt1*, as an initial step we described a few interesting cases of lineage-specific amino acid substitutions in Ranidea, ancestral branch specific substitution, and finally positive selection in one mutation in *H. sarda* and two *Dmrt1* duplications in *X.laevis* and *Pelodytes ibericus*.

Chromosome-level synteny is well preserved across anuran genomes, which is consistent with previous conclusions^94,95^. With the karyotype number varying between 10 and 14 across the 12 chromosome-level anuran genome assemblies, we detected several cases of chromosome fissions and fusions. However, genomic regions remain largely collinear at and surrounding the fission and fusion regions. For instance, between the genomes of *Eleutherodactylus coqui* (N=10) and *Engystomops pustulosus* (N=11), 5 chromosomal fusions and 4 additional fissions were detected. Yet, all these surrounding regions remain largely colinear between the species. The evidence of conserved synteny across anurans despite chromosomal fusions and fissions suggests conserved chromosomal evolution shaping the evolution of recombination, gene order and crossability in anurans, which is also consistent with the slow genome evolution found recently^95^. Indeed, extreme heterochiasmy, where recombination is restricted to telomeric regions in males but distributes evenly across female genome, has been documented in various anuran lineages with both XY and ZW sex chromosome systems (reviewed in ^39^). Furthermore, the conserved synteny could also be a consequence of rapid sex chromosome turnover, where sex chromosomes frequently alternate among related frog species and across lineages ^30,40^. Turnovers cause the whole genomic regions to constantly reshuffle and recombination restores regularly, which may prevent the large structural variation and genome rearrangement spread and get fixed across anuran genomes.

New master sex-determining genes can evolve by gene duplication or allelic diversification^96^. The only known master sex-determining gene in anurans, *DmW*, evolved via gene duplication of *Dmrt1* in the African clawed frog *X. laevis* ^47^. Rare duplications were detected in the ‘usual suspect’ master sex-determining genes, *Dmrt1*, *Foxl2, Sox3,* and *Bod1l* across 53 anuran genomes. In two species with a near-complete *Dmrt1* gene duplication (*Pelodvtes ibericus* and *Dendrobates tinctorius*) the master sex-determining gene remains unknown, and further research is needed to determine whether the duplicated copy acquired a master sex-determining role. Our analysis suggests that new master sex-determining genes rarely evolve by gene duplication in anurans, the exception being *X. laevis*. Allelic diversification seems like a more likely mechanism for the evolution of novel genetic sex determination in anurans. One caveat of our analysis is that due to the difficulty in Y chromosome assembly, most chromosome-level genome assemblies are from the homomorphic sex (either XX female or ZZ male) (Table S1), we therefore cannot evaluate if the duplications occurred on the Y or W chromosome.

New master sex-determining genes have been found to evolve via allelic diversification in various teleost fish lineages, including the fighting fish (*Betta splendens*), sablefish (*Anoplopoma fimbria*), three Scatophagidae fish species, and the black carp (*Mylopharyngodon piceus*) ^97–99^. In all these cases, different types of transposable elements were involved, and inserted in the promoter, or intronic region on the *Dmrt1* Y copy to directly or indirectly increase *Dmrt1* Y copy expression. The only case of allelic diversification in anura is the *Bod1l* gene in *Bufo virulis*^54^. It would be quite valuable to evaluate how common the allelic diversification mechanism is across anurans. This is not an easy task, as it would require phased X/Z and Y/W chromosome assemblies to identify the structural differences between the X/Z and Y/W copy.

Most anurans have highly conserved micro synteny on the same locations across homologous chromosomes, with occasional local inversions, such as in *Foxl2, Sox3* and *Bod1l*. Thus no gene translocation was detected for the four top candidate sex-determining genes (*Dmrt1*, *Bod1l*, *Foxl2* and *Sox3*) across various anuran genomes. Sex chromosome turnovers have been documented in many anurans, fishes, reptiles, insects and flowering plants ^27,30–35,40^, which can be driven by master sex-determining gene translocation, or novel mutations of genes involved in the sexual developmental pathway but took over a master sex-determining role ^30^. Despite the wide spread of sex chromosome turnovers across animals and flowering plants, the underlying genetic mechanism is largely unknown. The only identified lineages are Takifugu fishes, salmonid fishes and strawberries, where master sex-determining gene from one species (repeatedly) translocated to another chromosome to determine sex, often involving certain types of transposable elements ^33,35,37^. We did not detect any gene translocation in the 4 genes investigated here, suggesting gene translocation is not responsible for the rapid sex chromosome turnover across in Ranidae, as well as across anurans. Therefore, novel mutations on genes involved in the sexual developmental pathway is a more probable mechanism and would need to test with comparative genomics, requiring whole genome assembly to identify the sex-determining regions and candidate genes.

Theory predicts that at least at the initial phase, a newly evolved maser sex-determining gene is under somewhat positive selection and can sweep rapidly, if it counterbalances a suboptimal sex ratio or resolves a sex-linked conflict ^100,101^ In line with this prediction, both newly evolved master sex-determining genes *DmY* in medaka fish (*Oryzias latipes*) and *DmW* in the African clawed frogs (*X. laevis*) were under positive selection^58^. However, across 53 anurans, *Dmrt1* in the vast majority of them is under strong purifying selection, with very small values for both *d_N_* and d_S_. The strong purifying selection is also happening for those species where *Dmrt1* was identified as candidate master sex-determining gene or was located on the sex chromosomes. Together this suggests *Dmrt1* is under strong evolutionary constraints and mutations cannot accumulate on this gene. This is consistent with *Dmrt1*’s conserved role in the testis formation and development across various vertebrate groups ^23^. Furthermore, pleiotropic effects can also contribute to its evolutionary constraint. *Dmrt1* was reported to be a molecular controller for meiosis entry, and also a requirement for ovary development in the rabbit or certain fish ^91^. Interestingly, the only two cases where genes tended towards positive selection and more mutations contributed to *d_N_* than *d_S_* were the duplicated copy of *Dmrt1* in *X. laevis* and *Pelodytes ibericus*. In the case of *DmW* in *X. laevis*, the original copy *Dmrt1* is still functional and required for testis formation and development, yet the duplicated copy *DmW* was free from selection constraint and was under positive selection. Although the role of the duplicated copy of *Dmrt1* in *Pelodvtes ibericus* is unclear, a similar scenario where a duplicated copy freed itself from selection constraints and was under positive selection is possible.

The DM domain is a very conserved DNA-binding motif, shared by in total 9 *Dmrt* gene families across vertebrates lineages ^23^, of which three of them (*Dmrt7, Dmrt8, Dmrt2b*) are either mammal- or fish-specific. Furthermore, this domain is homologous to *Doublesex (dsx)* in insects, which regulates sexual differentiation ^102,103^. In mice, *Dmrt1* is necessary for testis maintenance and is sufficient to induce female-to-male cell fate reprogramming *in vivo*. *Dmrt1* expression in female ovary cells at around the sex-determining period could suppress expression *Foxl2* expression which is on the top of the female-determining pathway. Overall, *Dmrt1* is in the crossroad of sex determination, acting as transcriptional factor, interacting with various other downstream genes to determine male or female pathway ^97^. We detected several lineage-specific amino acid mutations combination on the DM domain. One of the most striking ones is the Ranidae-specific M-S-L-E mutations combination occurring at the positions of *Dmrt1* protein 30, 34, 44 and 56. Selection analysis showed these mutations were not under positive selection, but they could well be related to enhancing DNA binding affinity. Similarly, for *DmW* copy, H-T-I-Q-T-I-Q mutations were detected in the same DM protein region, most of them were not under positive selection, and the mutations are overall related to increase DNA binding affinity ^60^. Another amino acid substitution is the position 42 in the most ancestral branch, where they harboured the amino acid A, yet in the derived lineages with S. The current parsimonious scenarios both required two substitution change events, and it is unclear how the substitution changes, as well as the functional explanation remains unclear. The next step would be to evaluate the amino acid mutations in a molecular setting to understand whether this relates to DNA binding affinity, or link the mutations with broader life history traits and changes to find possible association for the function.

## 5. Conclusions

To conclude, chromosome-level synteny is highly conserved across anuran genomes, and four candidate master sex-determining genes showed rare gene duplications or translocations. Furthermore, gene translocation is unlikely driving the frequent sex chromosome turnovers across the anuran lineages. We therefore propose that new master sex-determining genes affecting the sexual development pathway most likely evolve in anurans via allelic diversification, and the next step is to evaluate this mechanism across anurans. This would require i) good chromosome-level genome assemblies to identify the sex-determining region, and ii) phased X/Z and Y/W chromosome assemblies to detect the diversification and novel mutation of the master sex-determining gene. All candidate sex-determining genes are under strong purifying selection, regardless of their role as master sex-determining gene or sex linkage. Gene duplication seems to have freed *Dmrt1* from selection constraints allowing it to assume a new adaptive function under positive selection. Finally, there are interesting lineage-specific amino acid mutations in Ranidea but their potential role in enhancing DNA binding affinity or other function requires further investigation.

## Supporting information

all supplementary figures, tables and file.

## Supplementary Materials

### Supplementary figures

Figure S1: Phylogenetic tree of the DM domain from all *DMRT* genes across anurans. The tree illustrates the evolutionary relationships among DM domains. Red-colored tips indicate partial duplications of the *DMRT1* DM domain.

Figure S2: RNA-seq reads aligned to exons 1 to 5 of *Dmrt1* in *Pelobates cultripes* (Western spadefoot toad). Grey and white boxes represent reads aligned to the genome, showing coverage across the exons. All exons displayed very low expression, with exon 1 showing minute read coverage. The RNA-seq dataset, derived from whole-body tadpole samples (SRR7817206, SRR7817207, SRR7817208, SRR7817209, SRR7817210, SRR7817211, SRR7817212, SRR7817213, SRR7817214, SRR7817215, SRR7817219, and SRR7817220), was mapped to the reference genome (GCA_933207985.1_aPelCul1.1_chrom_genomic.fna). The locations of each exon are shown as BED records in the bottom row.

Figure S3: RNA-seq reads aligned to the duplicated first copy of partial exon 1 and complete exons 3 to 5 of *Dmrt1* in *Pelobates cultripes* (Western spadefoot toad). Grey and white boxes represent reads aligned to the genome, showing coverage across the duplicated exons. All exons showed no expression. The RNA-seq dataset, derived from whole-body tadpole samples (SRR7817206, SRR7817207, SRR7817208, SRR7817209, SRR7817210, SRR7817211, SRR7817212, SRR7817213, SRR7817214, SRR7817215, SRR7817219, and SRR7817220), was mapped to the reference genome (*GCA_933207985.1_aPelCul1.1_chrom_genomic.fna*). The locations of each exon are shown as BED records in the bottom row.

Figure S4: RNA-seq reads aligned to the duplicated second copy of exons 3 to 5 of *Dmrt1* in *Pelobates cultripes* (Western spadefoot toad). Grey and white boxes represent reads aligned to the genome, showing coverage across the duplicated exons. All exons showed no expression. The RNA-seq dataset, derived from whole-body tadpole samples (SRR7817206, SRR7817207, SRR7817208, SRR7817209, SRR7817210, SRR7817211, SRR7817212, SRR7817213, SRR7817214, SRR7817215, SRR7817219, and SRR7817220), was mapped to the reference genome (*GCA_933207985.1_aPelCul1.1_chrom_genomic.fna*). The locations of each exon are shown as BED records in the bottom row.

Figure S5: RNA-seq reads aligned to *Dmrt1* in *Dendrobates tinctorius* (dyeing dart frog), showing exon-level coverage. All exons showed no read coverage. RNA-seq datasets from male brain and female liver, skin, and gut samples (SRR9304990, SRR17818134, SRR17818135, and SRR17818136) were mapped to the reference genome (*GCA_039654945.1_ASM3965494v1_genomic.fna*). The locations of each exon are shown as BED records in the bottom row.

Figure S6: RNA-seq reads aligned to the partially duplicated exons 1 and 3, and the completely duplicated exon 2 of *Dmrt1* in *Dendrobates tinctorius* (dyeing dart frog), showing exon-level coverage. All duplicated exons showed no read coverage. RNA-seq datasets from male brain and female liver, skin, and gut samples (SRR9304990, SRR17818134, SRR17818135, and SRR17818136) were mapped to the reference genome (*GCA_039654945.1_ASM3965494v1_genomic.fna*). The locations of each exon are shown as BED records in the bottom row.

Figure S7: RNA-seq reads aligned to *Dmrt1*, spanning exons 1 to 5 in *Rana temporaria* (Common frog). Grey boxes represent reads aligned to the genome, and thin lines connecting these boxes indicate spliced read alignments. RNA-seq datasets from male gonad samples (SRR8149028, SRR8149044, and SRR8149045) were mapped to the reference genome (*GCF_905171775.1_aRanTem1.1_genomic.fna*). The locations of each exon are shown as BED records in the top row.

Figure S8: RNA-seq reads aligned to the exon 2 duplication of *Dmrt1* in *Rana temporaria* (Common frog), with no reads supporting the duplicated region. RNA-seq datasets from male gonad samples (SRR8149028, SRR8149044, and SRR8149045) were mapped to the *Rana temporaria* reference genome (*GCF_905171775.1_aRanTem1.1_genomic.fna*). The duplicated region of exon 2 is indicated as a BED track in the top row.

Figure S9: RNA-seq reads aligned to exons 1 to 5 of *Dmrt1* in *Dendropsophus ebraccatus* (hourglass treefrog). Grey and white boxes represent reads aligned to the genome. The RNA-seq dataset from a female whole-body sample (SRR30172938) was mapped to the reference genome (*GCA_027789765.1_aDenEbr1.pat_genomic.fna*). The locations of each exon are shown as BED records in the bottom row.

Figure S10: RNA-seq reads aligned to Dmrt1 in *Ranitomeya imitator* (imitator poison frog), showing coverage across the exons. Only exon 5 displayed very low expression, while the other exons showed no read coverage. Grey and white boxes represent reads aligned to the genome. RNA-seq datasets from brain and skin samples (ERR3169416, SRR8275032, and SRR29319291) were mapped to the reference genome (GCA_032444005.1_aRanImi1.pri_genomic.fna). The locations of each exon are shown as BED records in the bottom row.

Figure S11: RNA-seq reads aligned to the duplicated exon 2 of *Dmrt1* in *Ranitomeya imitator* (imitator poison frog), showing no read coverage. RNA-seq datasets from brain and skin samples (ERR3169416, SRR8275032, and SRR29319291) were mapped to the reference genome (*GCA_032444005.1_aRanImi1.pri_genomic.fna*). The locations of each exon are shown as BED records in the bottom row.

Figure S12: *Dmrt1* expression based on read counts across 12 anurans. Gene expression levels were calculated using mapped reads obtained with HTSeq-count.

Figure S13: *Dmrt1* expression based on read counts across 2 anurans. Gene expression levels were calculated using mapped reads obtained with HTSeq-count.

Figure S14: dN, dS, and ω (dN/dS) for each species based on free-ratio models across 53 anuran species.

Figure S15: Plot of dS values of *Dmrt1* and its duplicated copies for 53 anurans, including the status of sex chromosomes reported in the literature. Boxplot of dS values.

Figure S16: Alignment of *Dmrt1* protein sequences (amino acid positions 1–191), visualized using Jalview.

Figure S17: Alignment of *Dmrt1* protein sequences (amino acid positions 190–370), visualized using Jalview.

### Supplementary tables

Table S1: Genome assemblies from NCBI for 54 amphibian species.

Table S2: Available raw genomic sequencing reads and the NCBI SRA number.

Table S3: RNAseq raw reads data from NCBI, and the related information on tissue, sex and developmental stage.

Table S4: Assess the completeness of genome/transcriptome assemblies using BUSCO Table S5: Exons of Dmrt1 and their lengths

Table S6: RNaseq data used for expression analysis between males and females, for species with at least one tissue in both sexes.

Table S7: The intact and duplicated copies of Dmrt1 in 8 anurans, along with their locations and lengths

Table S8: Dmrt1 gene synteny for annotated genomes of 12 species and BLAST hit synteny for chromosome-level genomes

Table S9: Foxl2 gene synteny for annotated genomes of 12 species

Table S10: Sox3 gene synteny for annotated genomes of 12 species and BLAST hit synteny for chromosome-level genomes

Table S11: Bod1l1 gene synteny for annotated genomes of 12 species and BLAST hit synteny for chromosome-level genomes

Table S12: (A) Omega, dN, and dS from the branch test for 54 species of Dmrt1 with DMW and Pelodytes_ibericus duplicated copy of Dmrt1 gene; (B) Omega, dN, and dS from the free-ratio model for 54 species of Dmrt1 with DMW and Pelodytes_ibericus duplicated copy of Dmrt1 gene; (C) Branch test for selection in 54 species of the Dmrt1 gene with DMW and Pelodytes_ibericus duplicated copy of Dmrt1 gene; (D) Branch-site test for 54 species of Dmrt1 with DMW and Pelodytes_ibericus duplicated copy of Dmrt1.

Table S13: (A) Omega, dN, and dS from the branch test for 52 species of Foxl2; (B) Branch test for selection in 52 species of the Foxl2

Table S14: (A) Omega, dN, and dS from the branch test for 53 species of Sox3 gene; (B) Branch test for selection in 53 species of the Sox3 gene

## Author Contributions

W-JM, SSS designed the study. SSS performed bioinformatic and related analyses. SSS, W-JM visualize the results. W-JM obtained the fundings for the project. W-JM, PV supervise the work. W-JM, SSS drafted the manuscript, which was improved and commented by PV. All authors agreed on the final version of the manuscript.

## Institutional Review Board Statement

Not applicable.

## Informed Consent Statement

Not applicable.

## Data Availability Statement

All analyzed data are downloaded from NCBI. The scripts for analyzing all related results are deposited in the Github repository and will make public upon paper acceptance: https://github.com/TheWMaLab/Duplication_evolution_MSDgenes_in_anurans

## Acknowledgements

We thank Kim Roelants for discussion and suggestions on selection analysis, and the Ma lab members for their discussion during the project development. The computations were performed at the VSC (Flemish Supercomputer Centre), funded by the Research Foundation - Flanders (FWO) and the Flemish Government.

