## Supplementary material for "Gene duplication, translocation and molecular evolution of *Dmrt1* and related sex-determining genes in anurans": all supplementary figures, tables and file.: Supplementary_figures_final.pdf

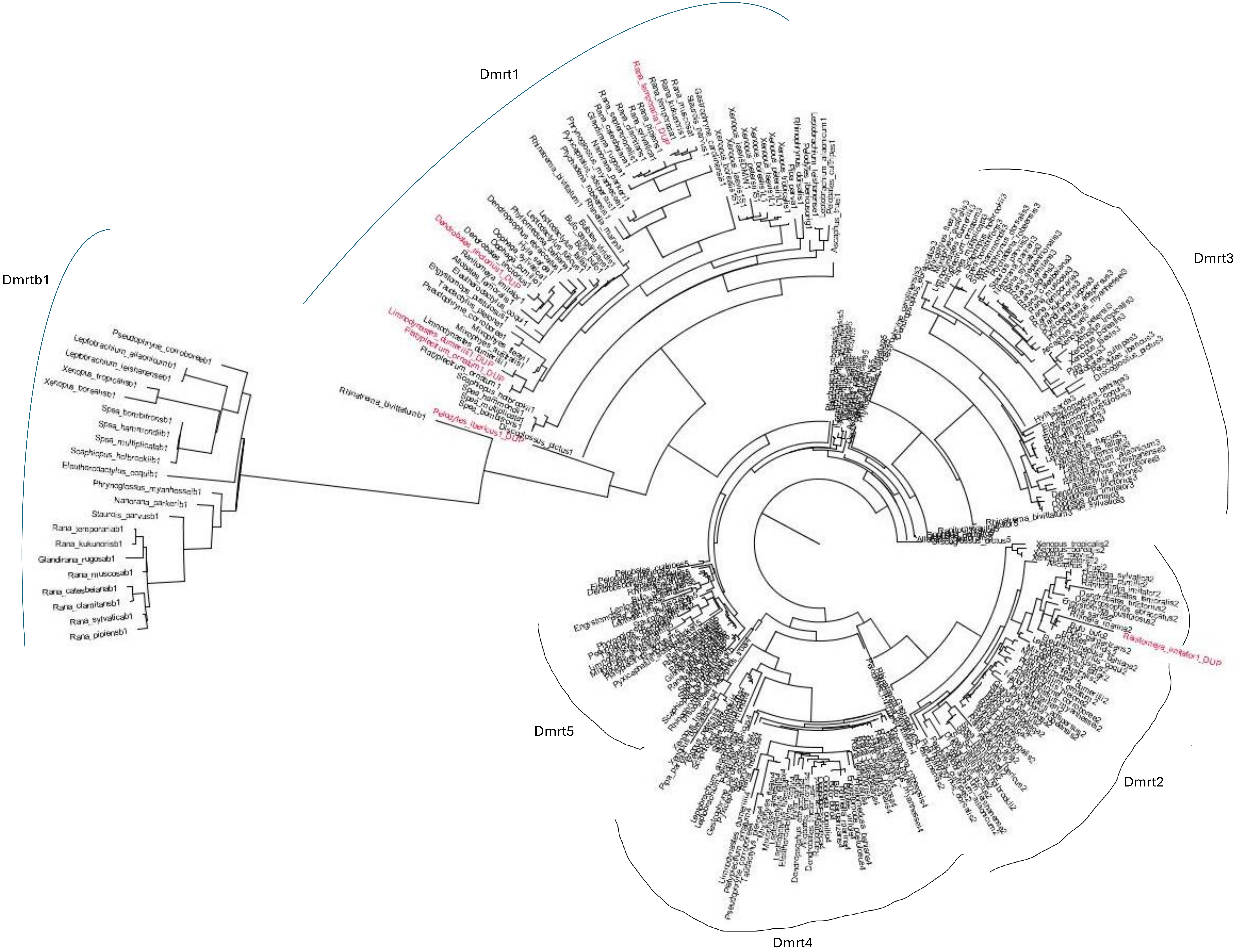


Figure S1

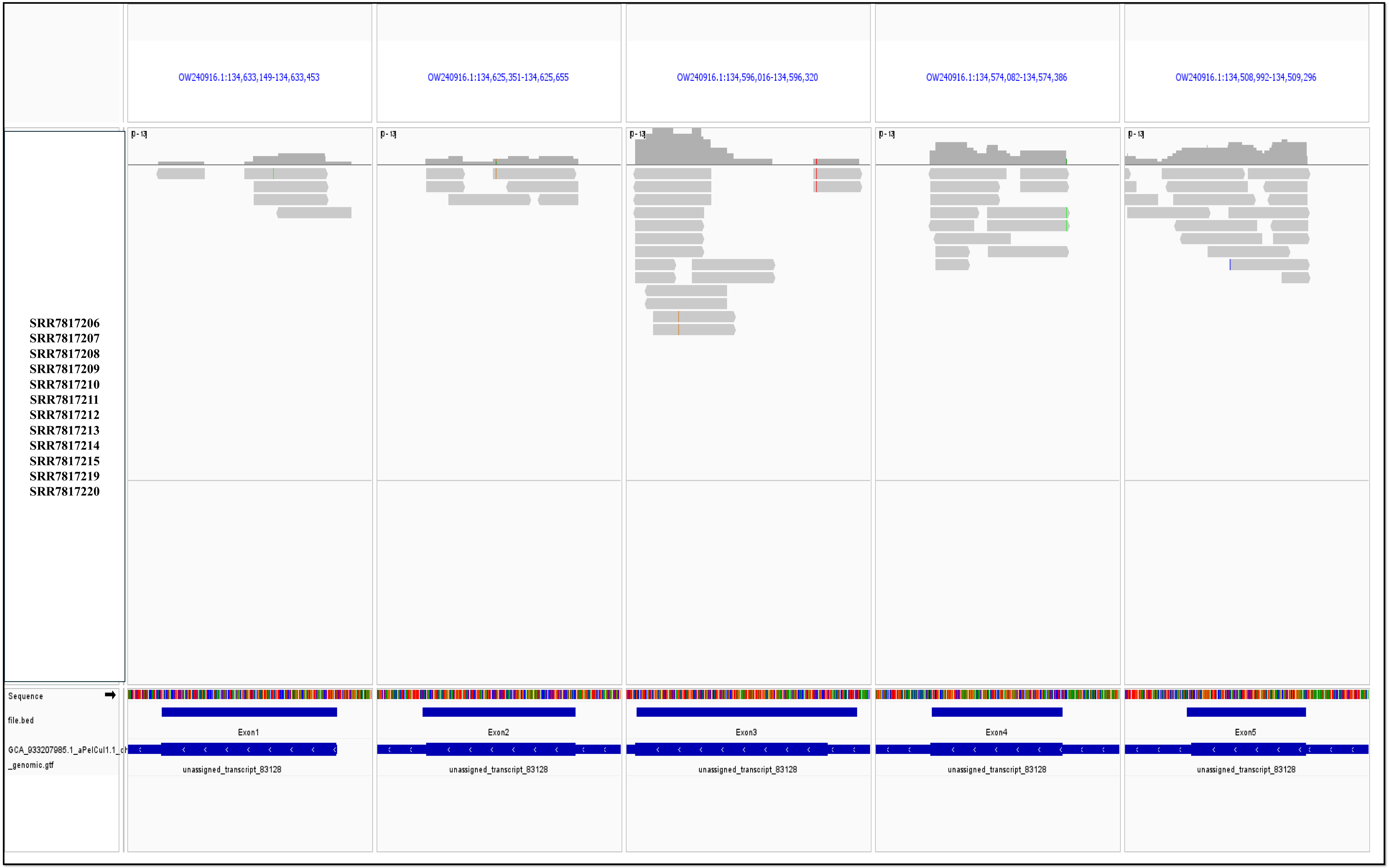
OW240916.1:134,633,149-134,633,453
OW240916.1:134,625,351-134,625,655
OW240916.1:134,596,016-134,596,320
OW240916.1:134,574,082-134,574,386
OW240916.1:134,508,992-134,509,296
SRR7817206
SRR7817207
SRR7817208
SRR7817209
SRR7817210
SRR7817211
SRR7817212
SRR7817213
SRR7817214
SRR7817215
SRR7817219
SRR7817220
Sequence
file.bed
GCA_933207985.1_aPelCul1.1_ch
_genomic.gtf
Exon1
Exon2
Exon3
Exon4
Exon5
unassigned_transcript_83128

Figure S2

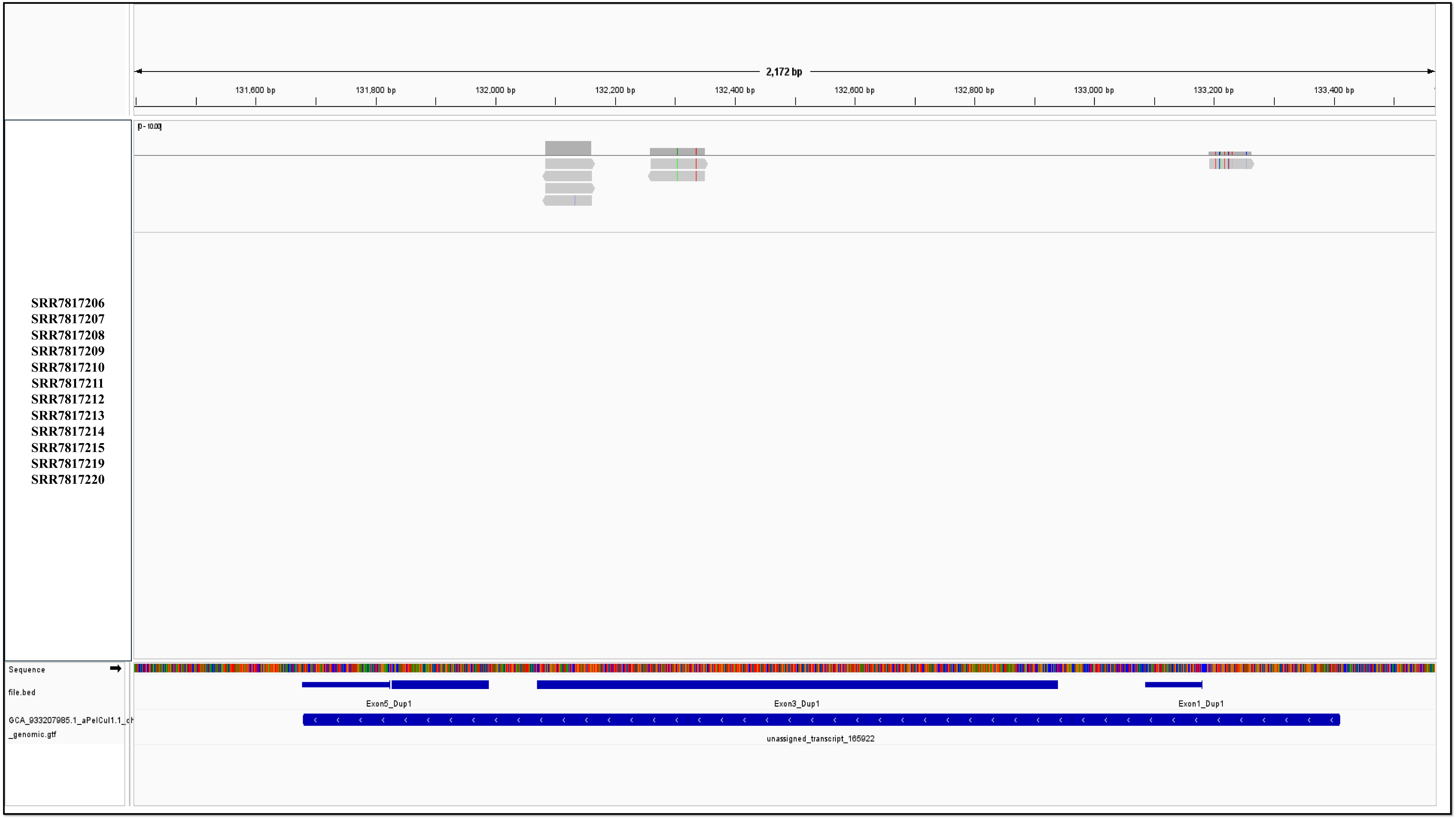
2,172 bp
131,600 bp
131,800 bp
132,000 bp
132,200 bp
132,400 bp
132,600 bp
132,800 bp
133,000 bp
133,200 bp
133,400 bp
SRR7817206
SRR7817207
SRR7817208
SRR7817209
SRR7817210
SRR7817211
SRR7817212
SRR7817213
SRR7817214
SRR7817215
SRR7817219
SRR7817220
Sequence
file.bed
Exon5_Dup1
Exon3_Dup1
Exon1_Dup1
unassigned_transcript_165922

Figure S3

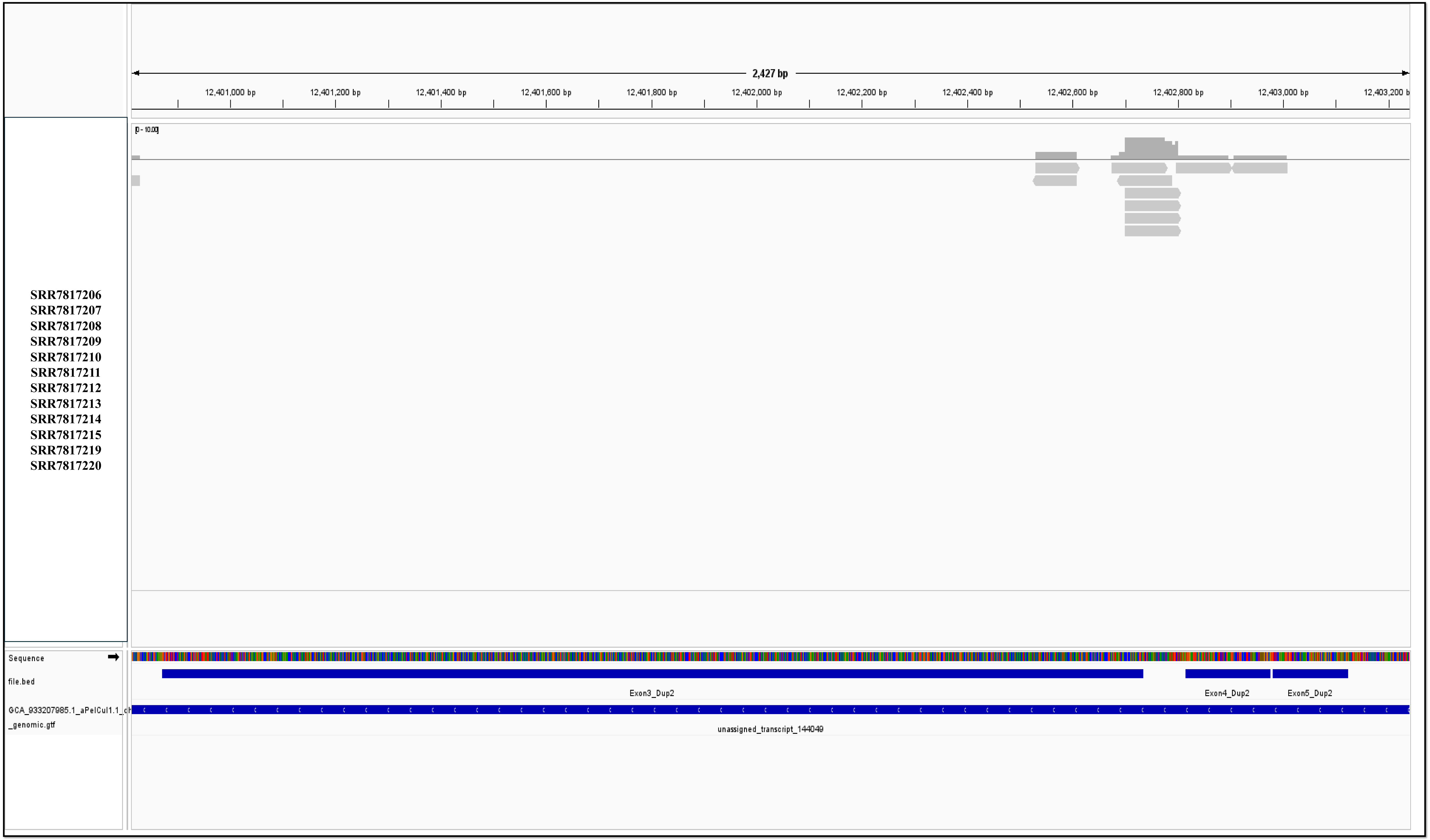
2,427 bp
12,401,000 bp
12,401,200 bp
12,401,400 bp
12,401,600 bp
12,401,800 bp
12,402,000 bp
12,402,200 bp
12,402,400 bp
12,402,600 bp
12,402,800 bp
12,403,000 bp
SRR7817206
SRR7817207
SRR7817208
SRR7817209
SRR7817210
SRR7817211
SRR7817212
SRR7817213
SRR7817214
SRR7817215
SRR7817219
SRR7817220
Sequence
file.bed
Exon3_Dup2
Exon4_Dup2
Exon5_Dup2
_genomic.gtf
unassigned_transcript_144049

Figure S4

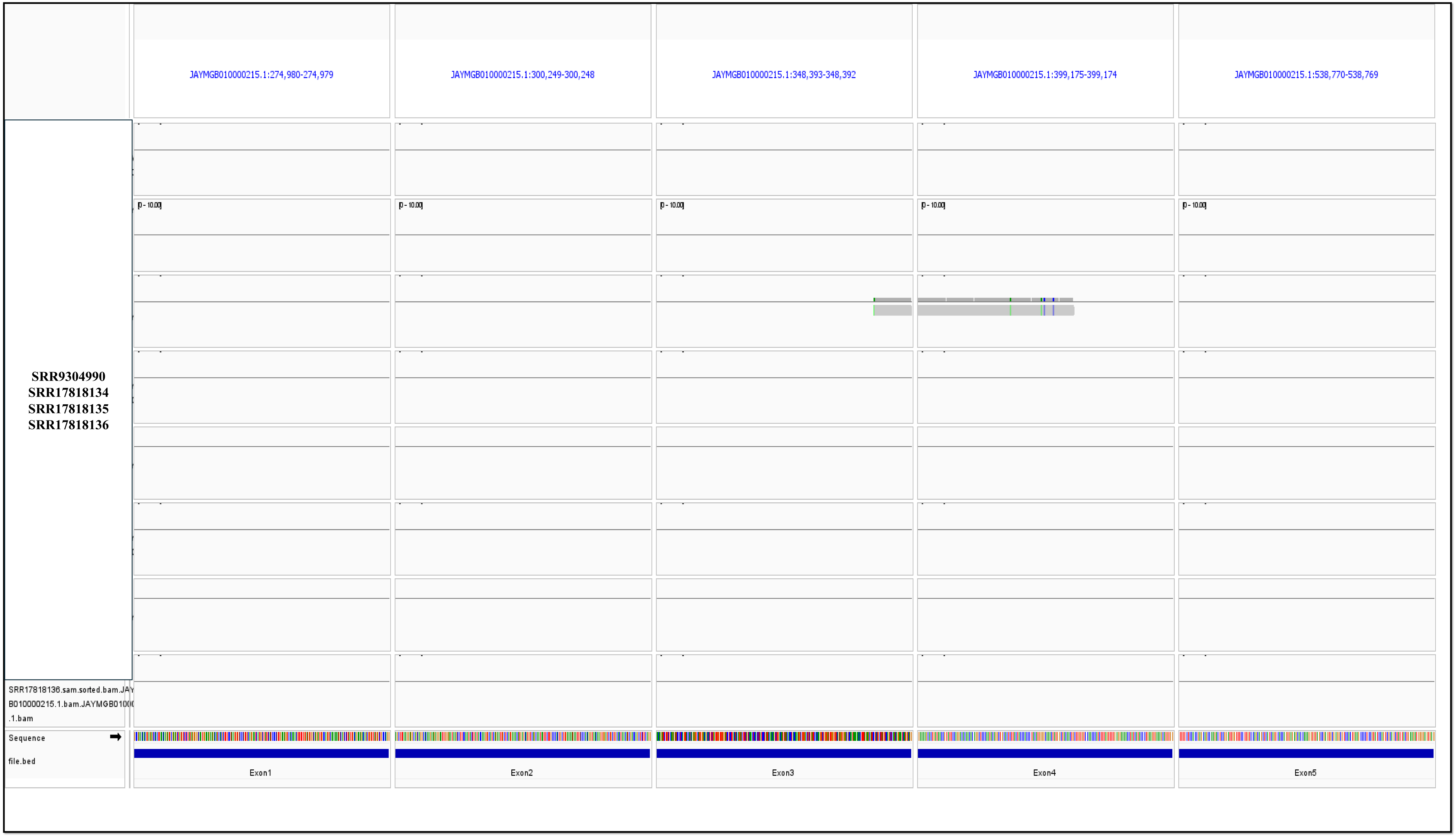
JAYMGB010000215.1:274,980-274,979
JAYMGB010000215.1:300,249-300,248
JAYMGB010000215.1:348,393-348,392
JAYMGB010000215.1:399,175-399,174
JAYMGB010000215.1:538,770-538,769
SRR9304990
SRR17818134
SRR17818135
SRR17818136
SRR17818136.sam.sorted.bam.JAY
B010000215.1.bam.JAYMGB0100
.1.bam
Sequence
file.bed
Exon1
Exon2
Exon3
Exon4
Exon5

Figure S5

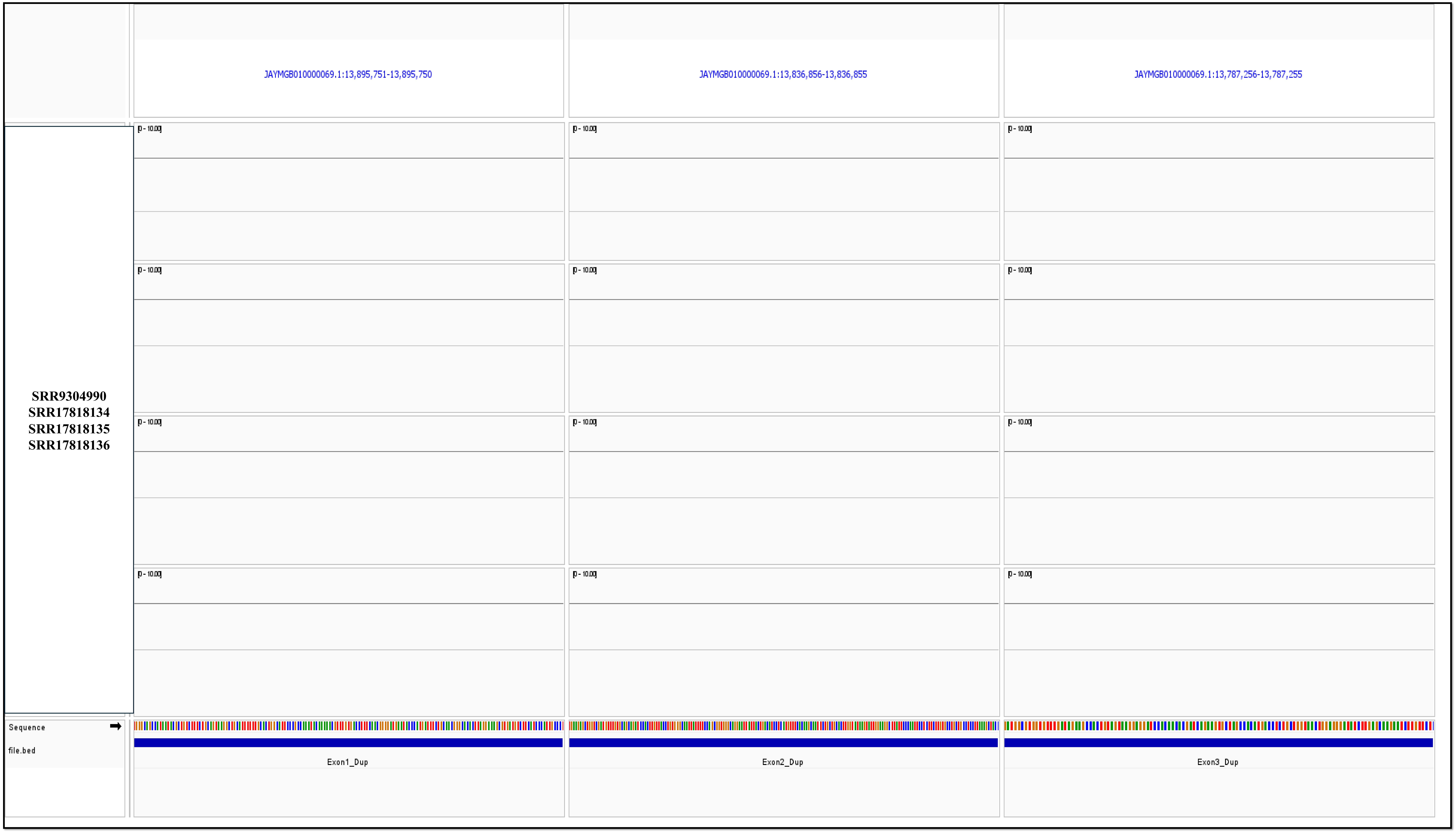

JAYMGB010000069.1:13,895,751-13,895,750
JAYMGB010000069.1:13,836,856-13,836,855
JAYMGB010000069.1:13,787,256-13,787,255
SRR9304990
SRR17818134
SRR17818135
SRR17818136
Sequence
file.bed
Exon1_Dup
Exon2_Dup
Exon3_Dup


Figure S6

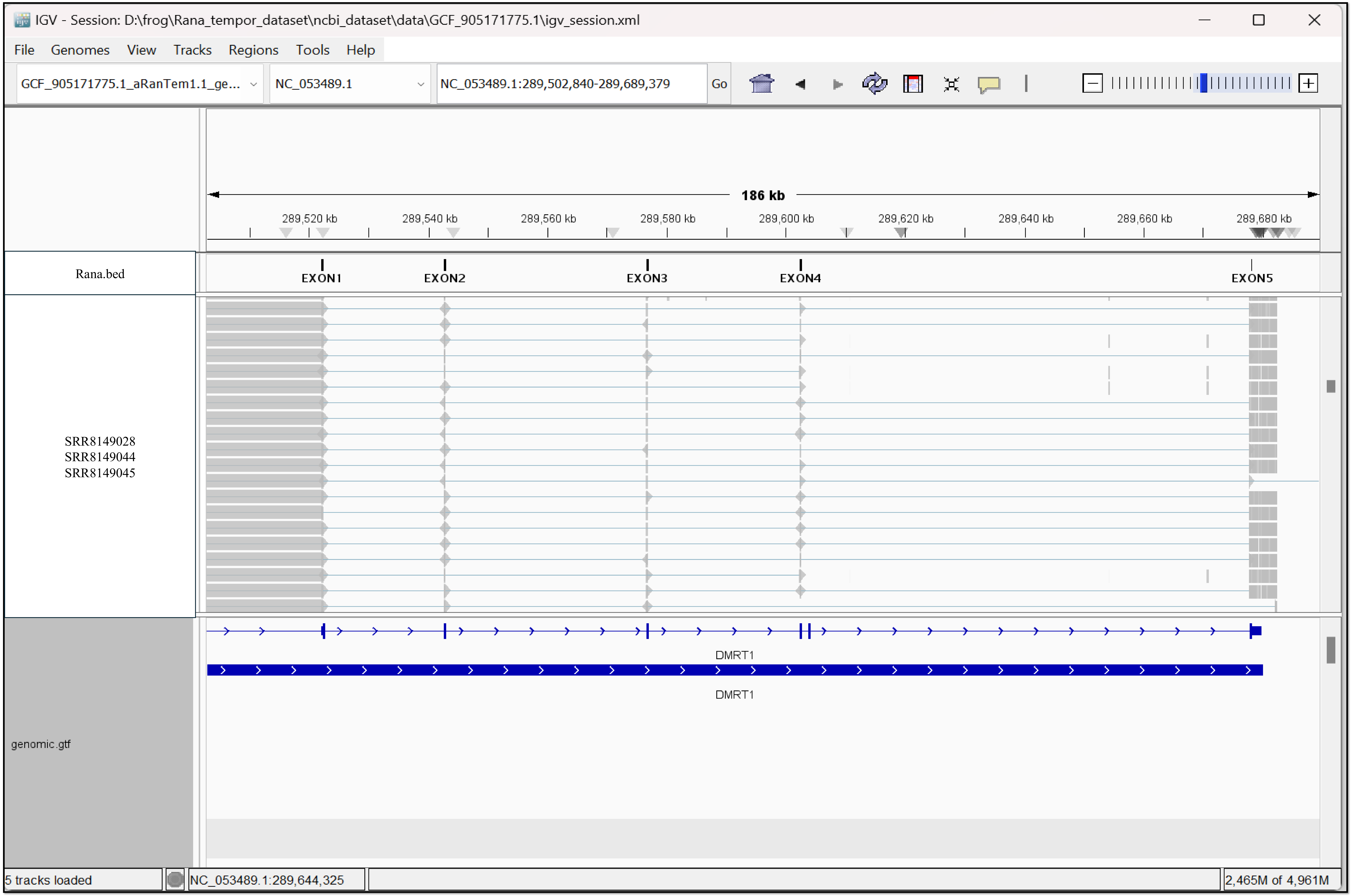
IGV - Session: D:\frog\Rana_tempor_dataset\ncbi_dataset\data\GCF_905171775.1\igv_session.xml
File Genomes View Tracks Regions Tools Help
GCF_905171775.1_aRanTem1.1_ge...
NC_053489.1
NC_053489.1:289,502,840-289,689,379
Go
186 kb
289,520 kb
289,540 kb
289,560 kb
289,580 kb
289,600 kb
289,620 kb
289,640 kb
289,660 kb
289,680 kb
Rana.bed
EXON1
EXON2
EXON3
EXON4
EXON5
SRR8149028
SRR8149044
SRR8149045
DMRT1
DMRT1
genomic.gtf
5 tracks loaded
NC_053489.1:289,644,325
2,465M of 4,961M

Figure S7

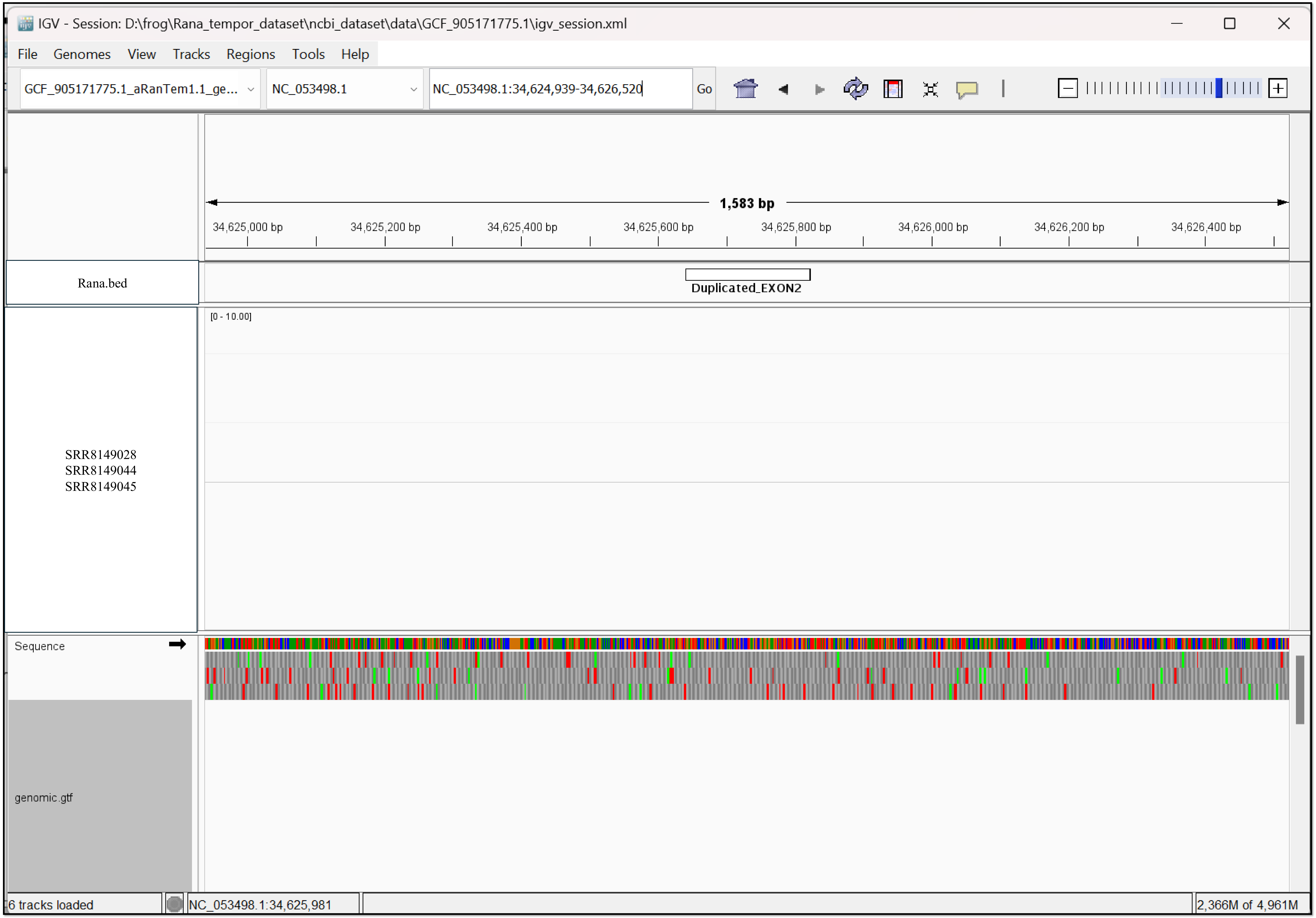
IGV - Session: D:\frog\Rana_tempor_dataset\ncbi_dataset\data\GCF_905171775.1\igv_session.xml
File
Genomes
View
Tracks
Regions
Tools
Help
GCF_905171775.1_aRanTem1.1_ge...
NC_053498.1
NC_053498.1:34,624,939-34,626,520
Go
1,583 bp
34,625,000 bp
34,625,200 bp
34,625,400 bp
34,625,600 bp
34,625,800 bp
34,626,000 bp
34,626,200 bp
34,626,400 bp
Rana.bed
Duplicated_EXON2
[0 - 10.00]
SRR8149028
SRR8149044
SRR8149045
Sequence
genomic.gtf
6 tracks loaded
NC_053498.1:34,625,981
2,366M of 4,961M

Figure S8

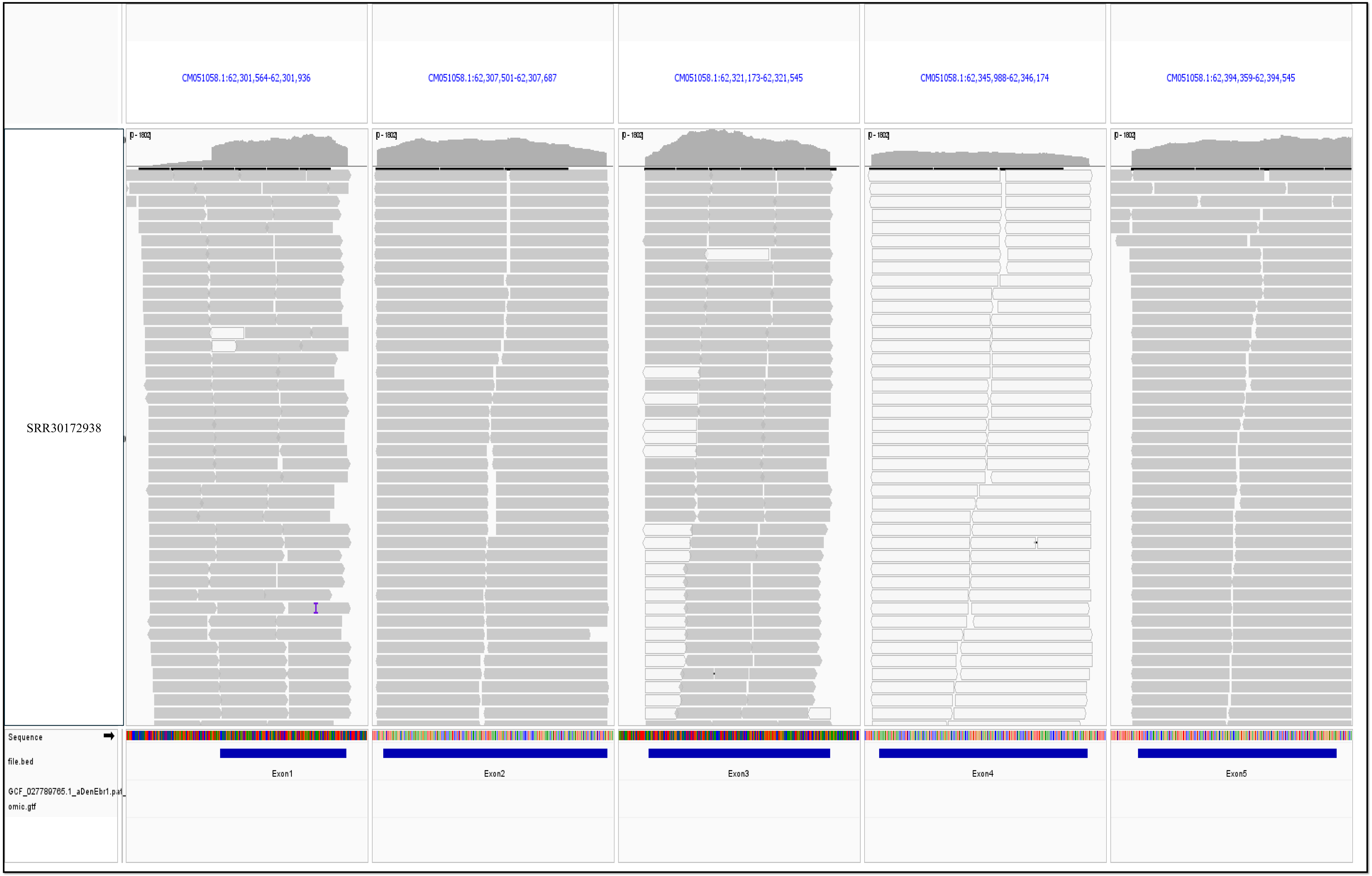
CM051058.1:62,301,564-62,301,936
CM051058.1:62,307,501-62,307,687
CM051058.1:62,321,173-62,321,545
CM051058.1:62,345,988-62,346,174
CM051058.1:62,394,359-62,394,545
SRR30172938
Sequence
file.bed
GCF_027789765.1_aDenEbr1.pat_
omic.gtf
Exon1
Exon2
Exon3
Exon4
Exon5

Figure S9

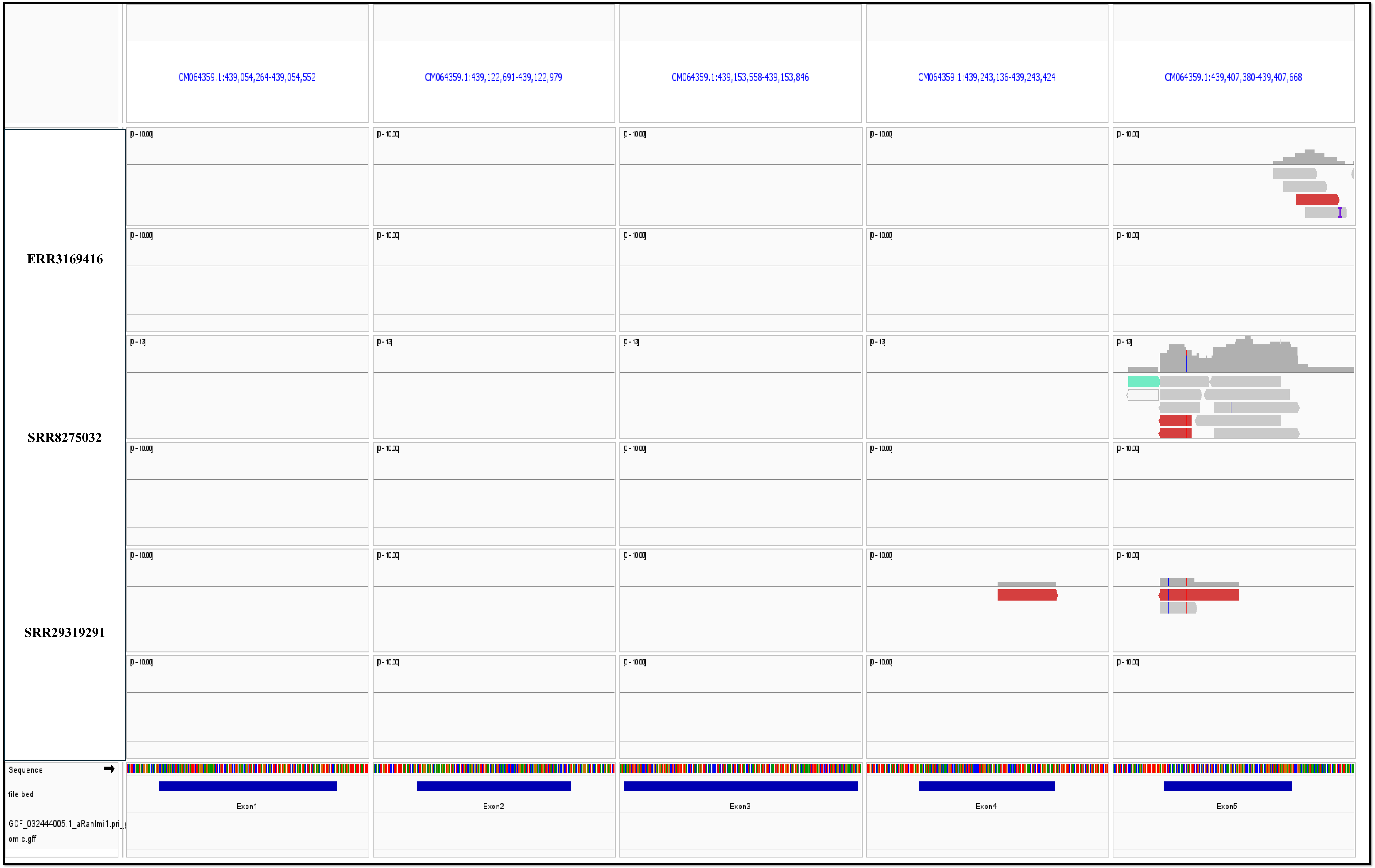
CM064359.1:439,054,264-439,054,552
CM064359.1:439,122,691-439,122,979
CM064359.1:439,153,558-439,153,846
CM064359.1:439,243,136-439,243,424
CM064359.1:439,407,380-439,407,668
ERR3169416
SRR8275032
SRR29319291
Sequence
file.bed
GCF_032444005.1_aRanImi1.pri_g
omic.gff
Exon1
Exon2
Exon3
Exon4
Exon5

Figure S10

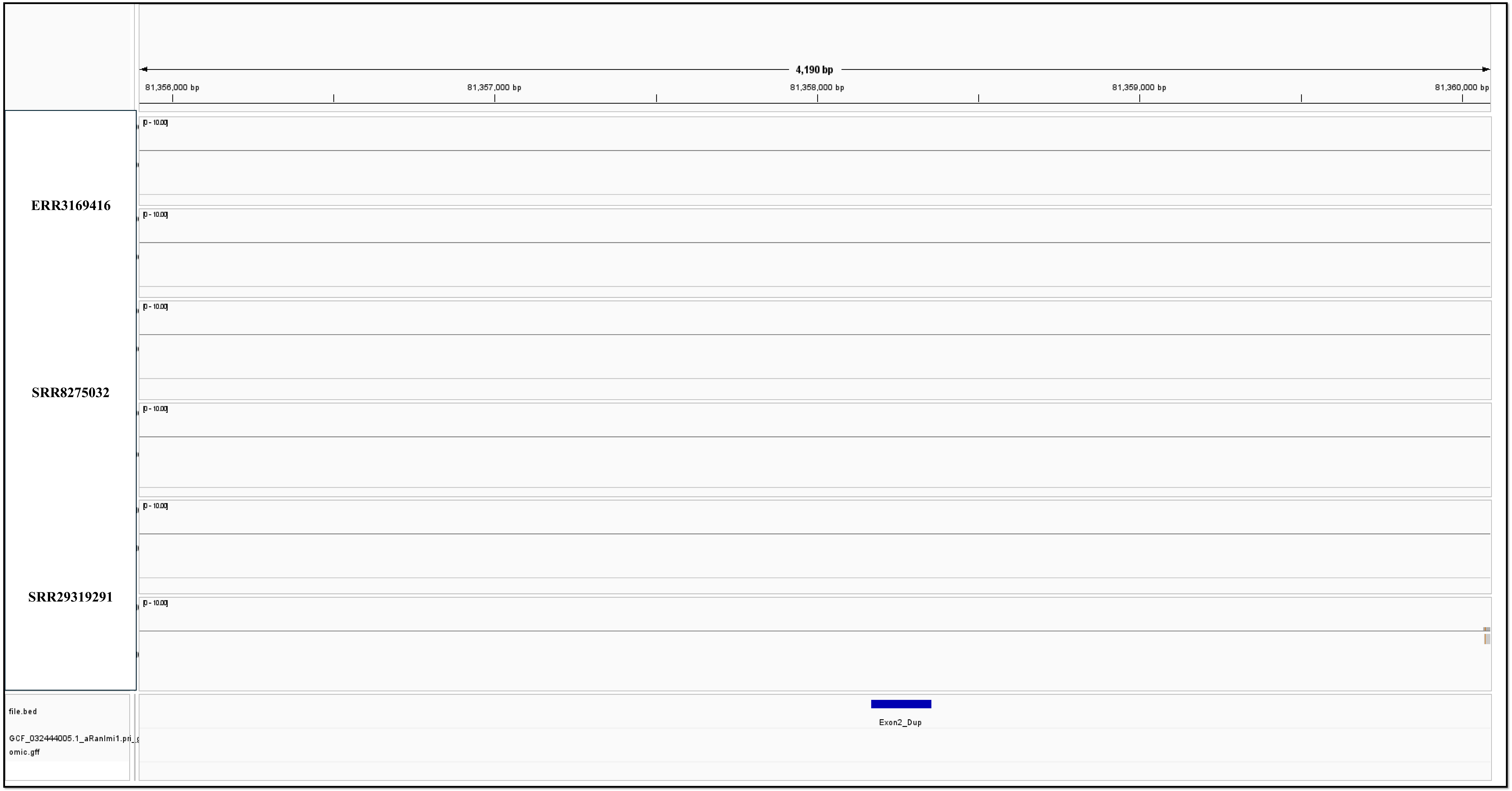
4,190 bp
81,356,000 bp
81,357,000 bp
81,358,000 bp
81,359,000 bp
81,360,000 bp
ERR3169416
SRR8275032
SRR29319291
file.bed
Exon2_Dup

Figure S11

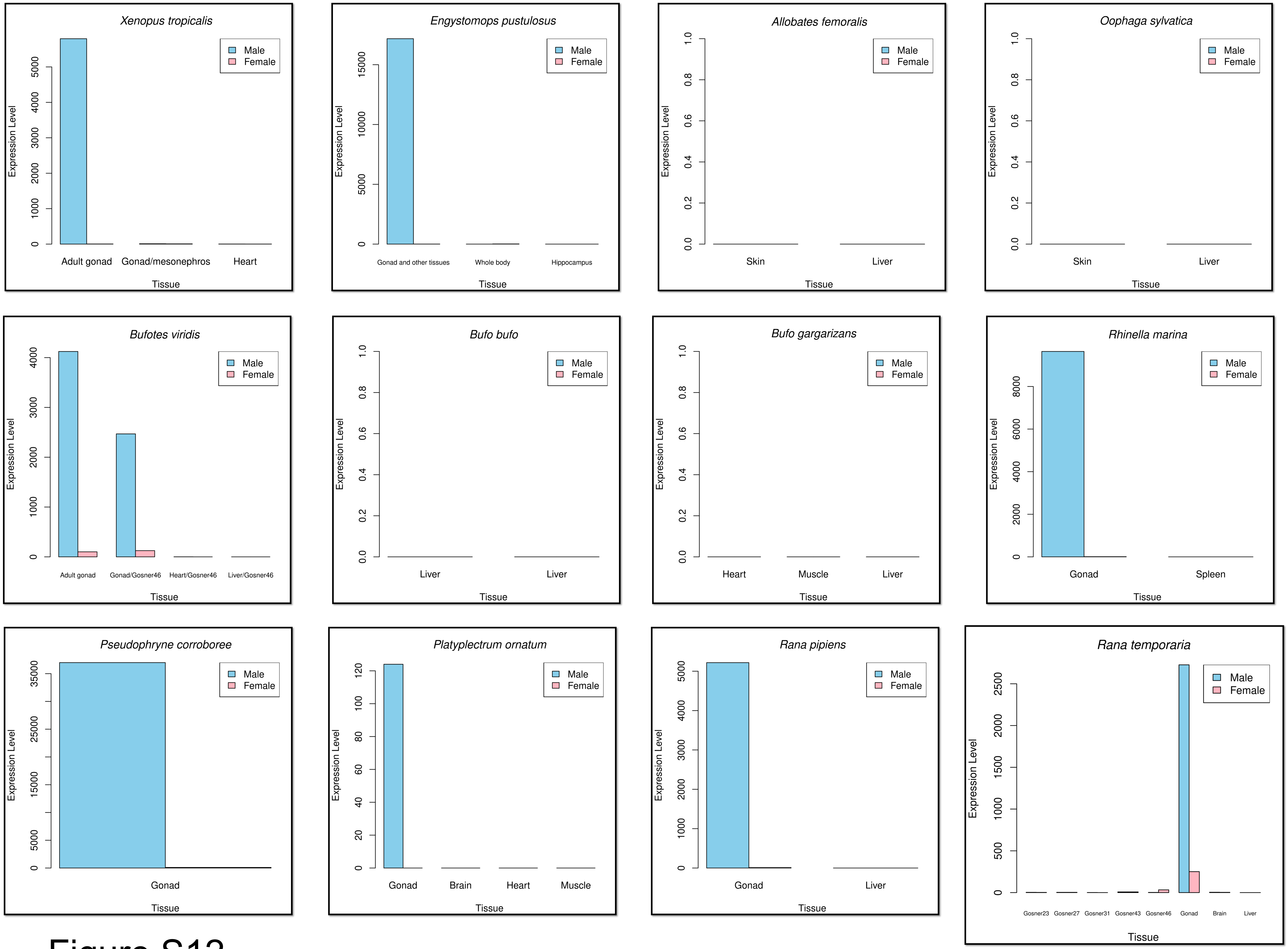

Xenopus tropicalis
Engystomops pustulosus
Allobates femoralis
Oophaga sylvatica
Bufotes viridis
Bufo bufo
Bufo gargarizans
Rhinella marina
Pseudophryne corroboree
Platyplectrum ornatum
Rana pipiens
Rana temporaria
Male
Female
Expression Level
Tissue
Adult gonad
Gonad/mesonephros
Heart
Gonad and other tissues
Whole body
Hippocampus
Skin
Liver
Gonad/Gosner46
Heart/Gosner46
Liver/Gosner46
Muscle
Gonad
Spleen
Brain
Gosner23
Gosner27
Gosner31
Gosner43
Gosner46


Figure S12

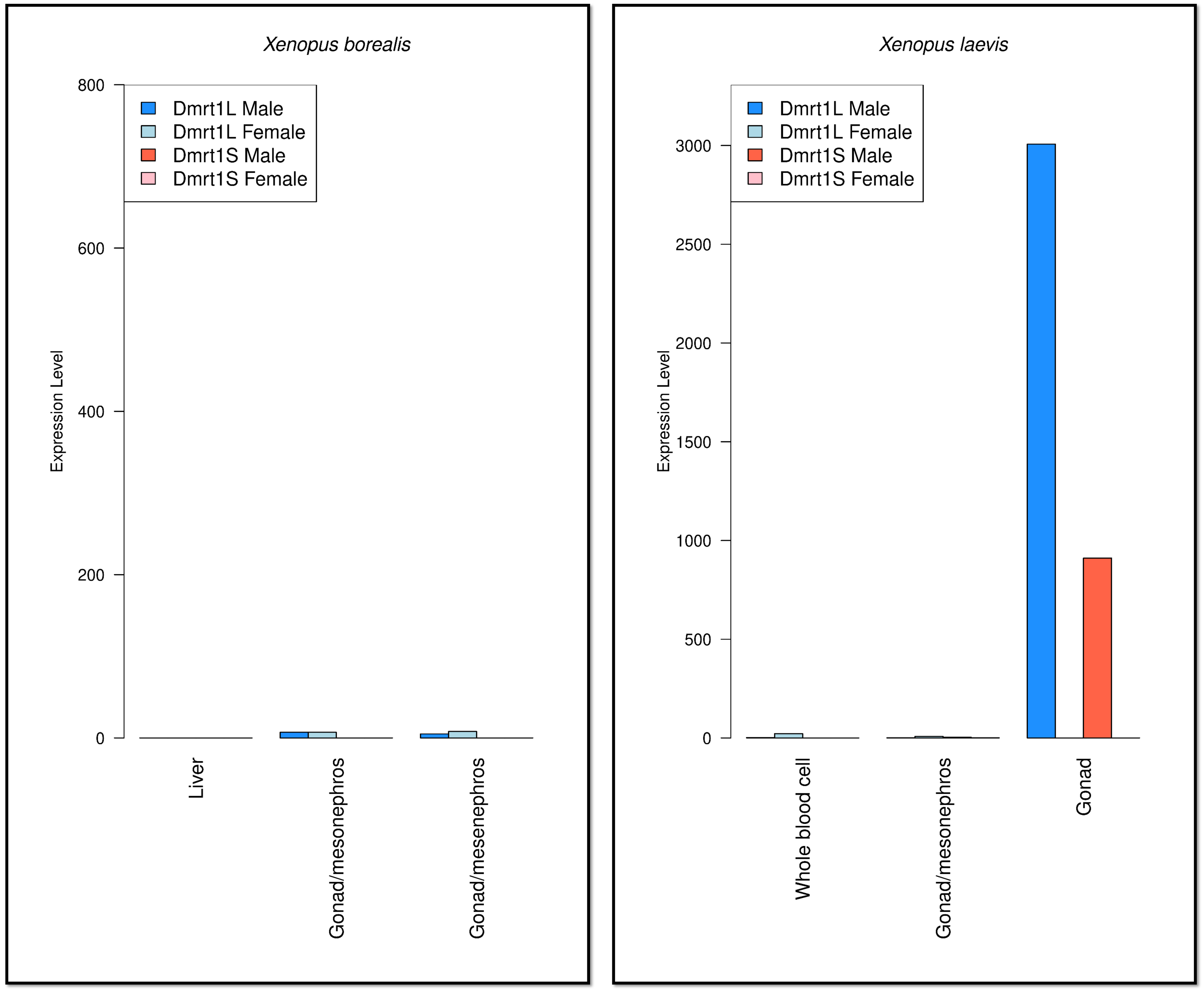

Xenopus borealis
800
600
400
200
0
Expression Level
Dmrt1L Male
Dmrt1L Female
Dmrt1S Male
Dmrt1S Female
Liver
Gonad/mesonephros
Gonad/mesenephros
Xenopus laevis
3000
2500
2000
1500
1000
500
0
Expression Level
Dmrt1L Male
Dmrt1L Female
Dmrt1S Male
Dmrt1S Female
Whole blood cell
Gonad/mesonephros
Gonad


Figure S13

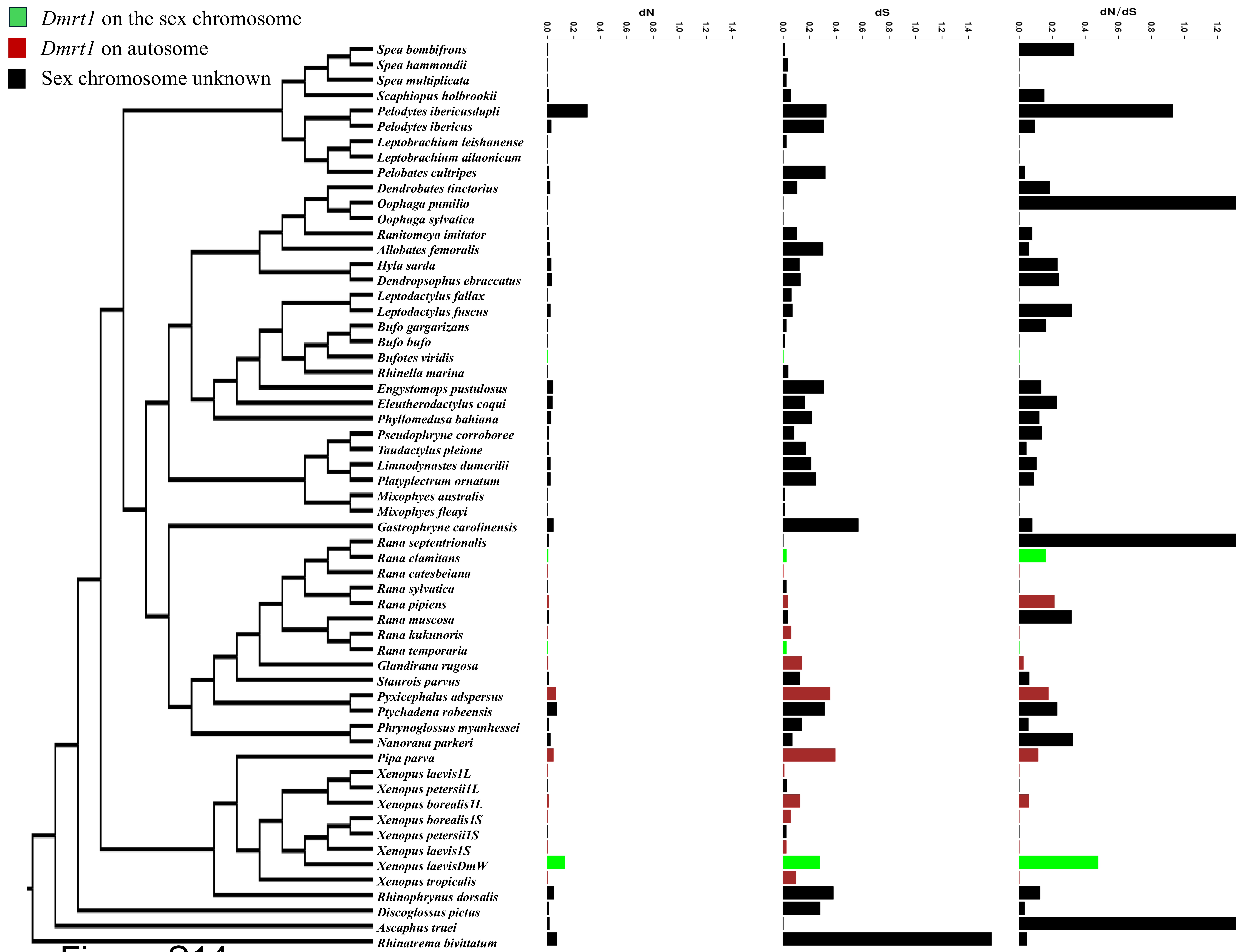
Dmrt1 on the sex chromosome
Dmrt1 on autosome
Sex chromosome unknown
dN
dS
dN/dS
0.0
0.2
0.4
0.6
0.8
1.0
1.2
1.4
Spea bombifrons
Spea hammondii
Spea multiplicata
Scaphiopus holbrookii
Pelodytes ibericusdupli
Pelodytes ibericus
Leptobrachium leishanense
Leptobrachium ailaonicum
Pelobates cultripes
Dendrobates tinctorius
Oophaga pumilio
Oophaga sylvatica
Ranitomeya imitator
Allobates femoralis
Hyla sarda
Dendropsophus ebraccatus
Leptodactylus fallax
Leptodactylus fuscus
Bufo gargarizans
Bufo bufo
Bufotes viridis
Rhinella marina
Engystomops pustulosus
Eleutherodactylus coqui
Phyllomedusa bahiana
Pseudophryne corroboree
Taudactylus pleione
Limnodynastes dumerilii
Platyplectrum ornatum
Mixophyes australis
Mixophyes fleayi
Gastrophryne carolinensis
Rana septentrionalis
Rana clamitans
Rana catesbeiana
Rana sylvatica
Rana pipiens
Rana muscosa
Rana kukunoris
Rana temporaria
Glandirana rugosa
Staurois parvus
Pyxicephalus adspersus
Ptychadena robeensis
Phrynoglossus myanhessei
Nanorana parkeri
Pipa parva
Xenopus laevis1L
Xenopus petersii1L
Xenopus borealis1L
Xenopus borealis1S
Xenopus petersii1S
Xenopus laevis1S
Xenopus laevisDmW
Xenopus tropicalis
Rhinophrynus dorsalis
Discoglossus pictus
Ascaphus truei
Rhinatrema bivittatum

Figure S14

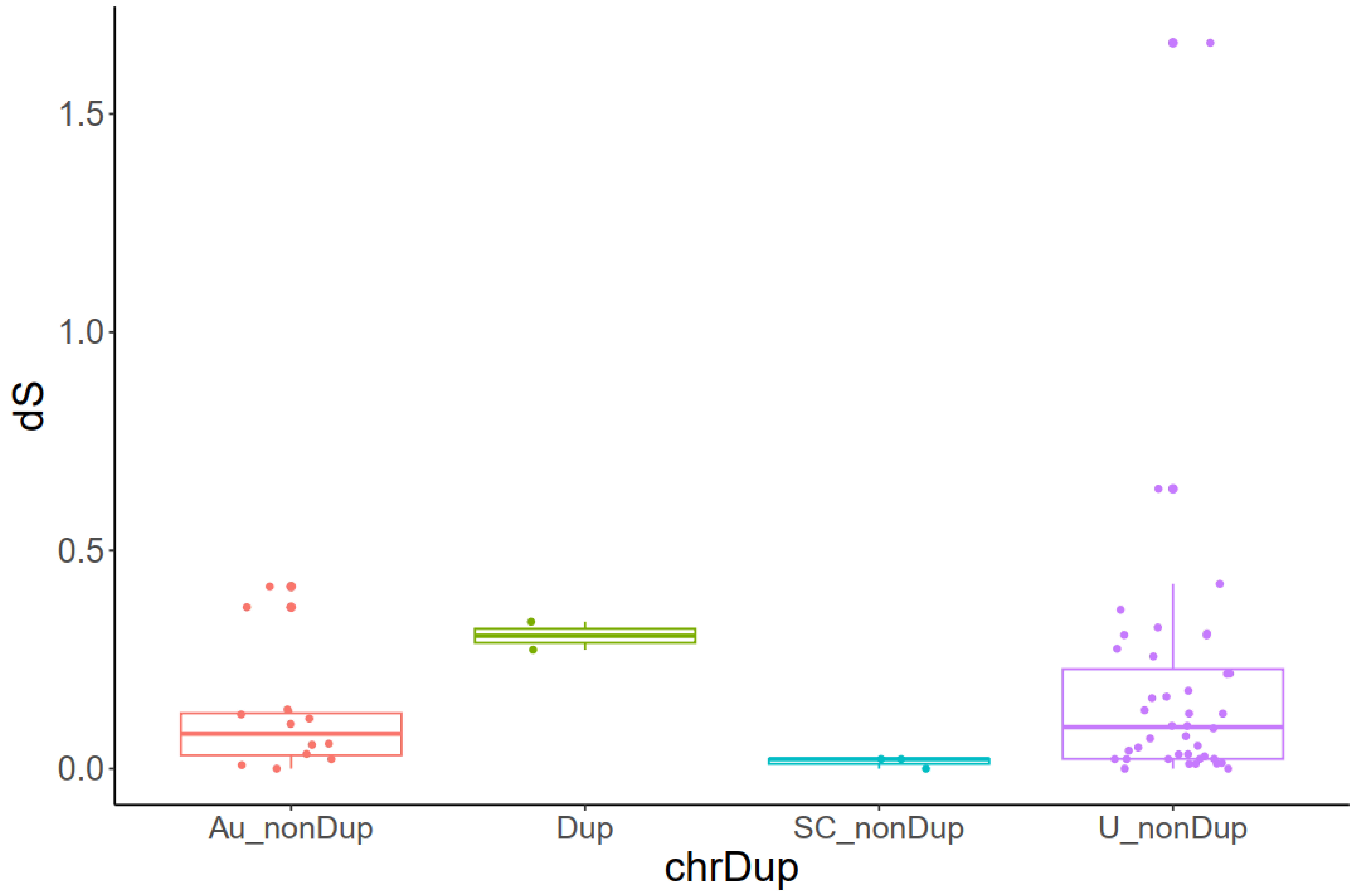


Figure S15

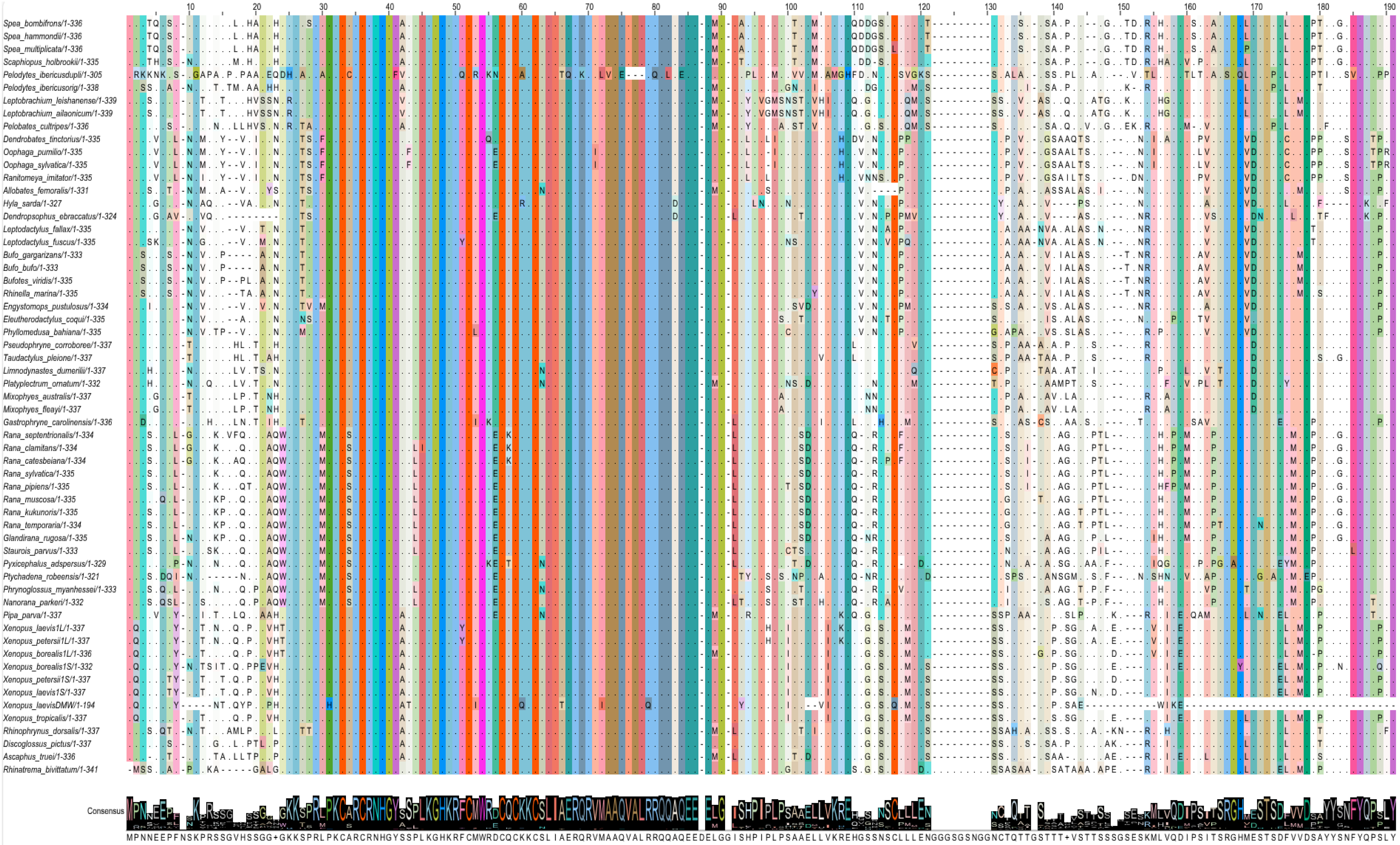
Spea_bombifrons/1-336
Spea_hammondii/1-336
Spea_multiplicata/1-336
Scaphiopus_holbrookii/1-335
Pelodytes_ibericusdupli/1-305
Pelodytes_ibericusorig/1-338
Leptobrachium_leishanense/1-339
Leptobrachium_ailaonicum/1-339
Pelobates_cultripes/1-336
Dendrobates_tinctorius/1-335
Oophaga_pumilio/1-335
Oophaga_sylvatica/1-335
Ranitomeya_imitator/1-335
Allobates_femoralis/1-331
Hyla_sarda/1-327
Dendropsophus_ebraccatus/1-324
Leptodactylus_fallax/1-335
Leptodactylus_fuscus/1-335
Bufo_gargarizans/1-333
Bufo_bufo/1-333
Bufotes_viridis/1-335
Rhinella_marina/1-335
Engystomops_pustulosus/1-334
Eleutherodactylus_coqui/1-335
Phyllomedusa_bahiana/1-335
Pseudophryne_corroboree/1-337
Taudactylus_pleione/1-337
Limnodynastes_dumerilii/1-337
Platyplectrum_ornatum/1-332
Mixophyes_australis/1-337
Mixophyes_fleayi/1-337
Gastrophryne_carolinensis/1-336
Rana_septentrionalis/1-334
Rana_clamitans/1-334
Rana_catesbeiana/1-334
Rana_sylvatica/1-335
Rana_pipiens/1-335
Rana_muscosa/1-335
Rana_kukunoris/1-335
Rana_temporaria/1-334
Glandirana_rugosa/1-335
Staurois_parvus/1-333
Pyxicephalus_adspersus/1-329
Ptychadena_robeensis/1-321
Phrynoglossus_myanhessei/1-333
Nanorana_parkeri/1-332
Pipa_parva/1-337
Xenopus_laevis1L/1-337
Xenopus_petersii1L/1-337
Xenopus_borealis1L/1-336
Xenopus_borealis1S/1-332
Xenopus_petersii1S/1-337
Xenopus_laevis1S/1-337
Xenopus_laevisDMW/1-194
Xenopus_tropicalis/1-337
Rhinophrynus_dorsalis/1-337
Discoglossus_pictus/1-337
Ascaphus_truei/1-336
Rhinatrema_bivittatum/1-341
Consensus
MPNNEEPFNSKPRSSGVHSSGG+GKKSPRLPKCARCRNHGYSSPLKGHKRFCMWRDCQCKKCSLIAERQRVMAAQVALRRQQAQEEDELGGISHPIPLPSAAELLVKREHGSSNSCLLLENGGGSGSNGGNCTQTTGSTTT+VSTTSSSGSESKMLVQDIPSITSRGHMESTSDFVVDSAYYSNFYQPSLY

Figure S16

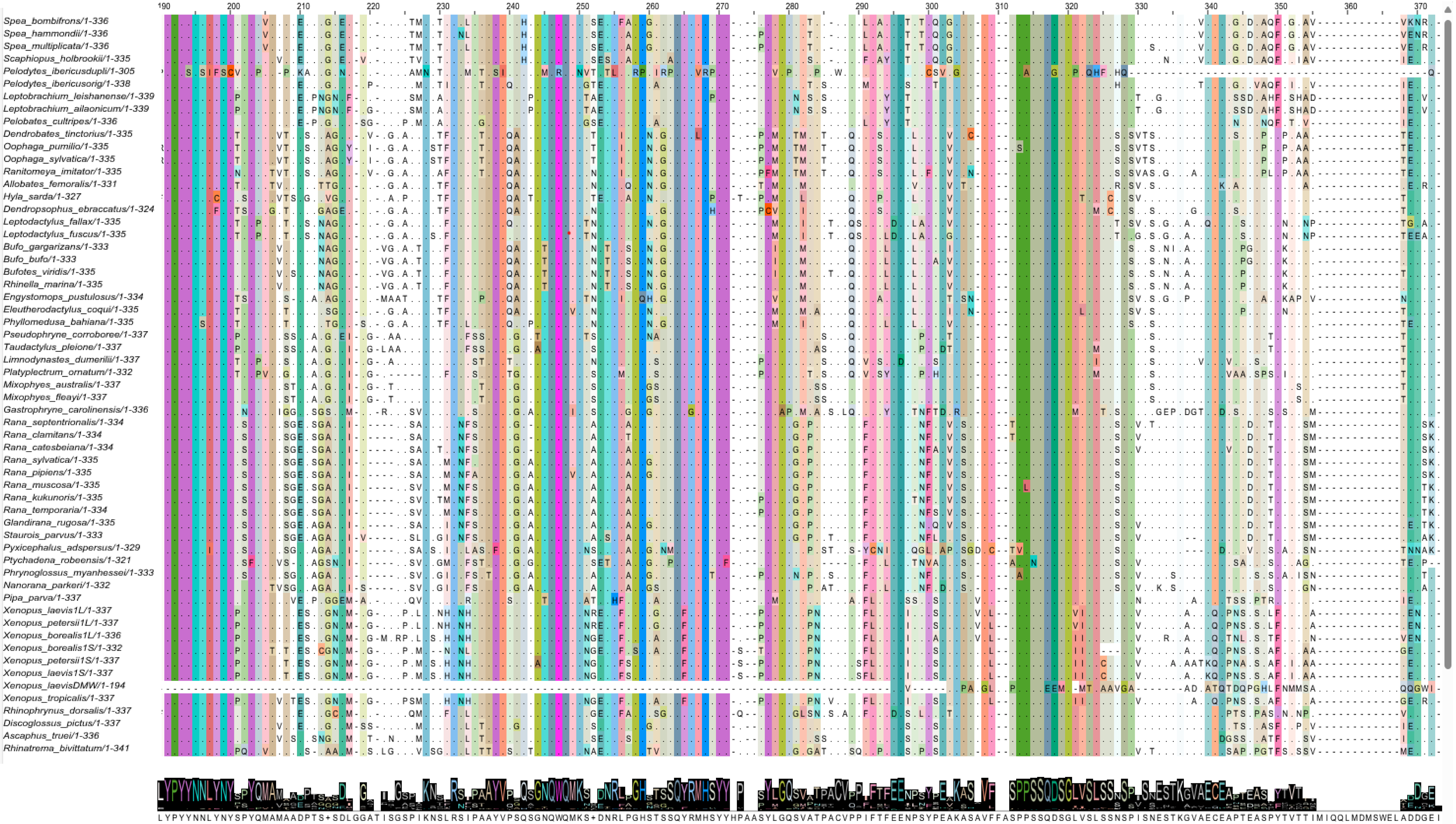
Spea_bombifrons/1-336
Spea_hammondii/1-336
Spea_multiplicata/1-336
Scaphiopus_holbrookii/1-335
Pelodytes_ibericusdupli/1-305
Pelodytes_ibericusorig/1-338
Leptobrachium_leishanense/1-339
Leptobrachium_ailaonicum/1-339
Pelobates_cultripes/1-336
Dendrobates_tinctorius/1-335
Oophaga_pumilio/1-335
Oophaga_sylvatica/1-335
Ranitomeya_imitator/1-335
Allobates_femoralis/1-331
Hyla_sarda/1-327
Dendropsophus_ebraccatus/1-324
Leptodactylus_fallax/1-335
Leptodactylus_fuscus/1-335
Bufo_gargarizans/1-333
Bufo_bufo/1-333
Bufotes_viridis/1-335
Rhinella_marina/1-335
Engystomops_pustulosus/1-334
Eleutherodactylus_coqui/1-335
Phyllomedusa_bahiana/1-335
Pseudophryne_corroboree/1-337
Taudactylus_pleione/1-337
Limnodynastes_dumerilii/1-337
Platyplectrum_ornatum/1-332
Mixophyes_australis/1-337
Mixophyes_fleayi/1-337
Gastrophryne_carolinensis/1-336
Rana_septentrionalis/1-334
Rana_clamitans/1-334
Rana_catesbeiana/1-334
Rana_sylvatica/1-335
Rana_pipiens/1-335
Rana_muscosa/1-335
Rana_kukunoris/1-335
Rana_temporaria/1-334
Glandirana_rugosa/1-335
Staurois_parvus/1-333
Pyxicephalus_adspersus/1-329
Ptychadena_robeensis/1-321
Phrynoglossus_myanhessei/1-333
Nanorana_parkeri/1-332
Pipa_parva/1-337
Xenopus_laevis1L/1-337
Xenopus_petersii1L/1-337
Xenopus_borealis1L/1-336
Xenopus_borealis1S/1-332
Xenopus_petersii1S/1-337
Xenopus_laevis1S/1-337
Xenopus_laevisDMW/1-194
Xenopus_tropicalis/1-337
Rhinophrynus_dorsalis/1-337
Discoglossus_pictus/1-337
Ascaphus_truei/1-336
Rhinatrema_bivittatum/1-341

Figure S17
